# The evolution of red colour vision is linked to coordinated rhodopsin tuning in lycaenid butterflies

**DOI:** 10.1101/2020.04.06.027102

**Authors:** Marjorie A. Liénard, Gary D. Bernard, Andrew A. Allen, Jean-Marc Lassance, Siliang Song, Richard Rabideau Childers, Nanfang Yu, Dajia Ye, Adriana Stephenson, Wendy A. Valencia-Montoya, Shayla Salzman, Melissa R.L. Whitaker, Michael Calonje, Feng Zhang, Naomi E. Pierce

**Affiliations:** The Broad Institute of MIT and Harvard, Cambridge, MA 02142, USA; Department of Organismic and Evolutionary Biology and Museum of Comparative Zoology, Harvard University, Cambridge, MA 02138, USA; Department of Electrical & Computer Engineering, University of Washington, Seattle, WA, USA; Department of Applied Physics and Applied Mathematics, Columbia University, New York, NY 10027, USA; Montgomery Botanical Center, 11901 Old Cutter Road, Miami, FL 33156, USA

**Keywords:** molecular evolution, ecological adaptation, visual system/vision, rhodopsin, spectral sensitivity, insects, Lepidoptera

## Abstract

Colour vision is largely mediated by changes in number, expression, and spectral properties of rhodopsins, but the genetic mechanisms underlying adaptive shifts in spectral sensitivity remain largely unexplored. Using *in vivo* photochemistry, optophysiology, and *in vitro* functional assays, we link variation in eye spectral sensitivity at long wavelengths to species-specific absorbance spectra for LW opsins in lycaenid butterflies. In addition to loci specifying an ancestral green-absorbing rhodopsin with maximum spectral sensitivity (λ_max_) at 520-530 nm in *Callophrys sheridanii* and *Celastrina ladon*, we find a novel form of red-shifted LW rhodopsin at λ_max_ = 565-570 nm in *Arhopala japonica* and *Eumaeus atala*. Furthermore, we show that *Ca. sheridanii* and *Ce. ladon* exhibit a smaller bathochromic shift at BRh2 (480-489 nm), and with the ancestral LW rhodopsin, cannot perceive visible red light beyond 600 nm. In contrast, molecular variation at the LW opsin in *A. japonica* and *E. atala* is coordinated with tuning of the blue opsin that also shifts sensitivity to longer wavelengths enabling colour discrimination up to 617 nm. We then use *E. atala* as a model to examine the interplay between red and blue spectral sensitivity. Owing to blue duplicate expression, the spatial distribution of opsin mRNAs within an ommatidium defines an expanded retinal stochastic mosaic of at least six opsin-based photoreceptor classes. Our mutagenesis *in vitro* assays with BRh1 (λ_max_ = 435 nm) chimeric blue rhodopsins reveal four main residues contributing to the 65 nm bathochromic shift towards BRh2 (λ_max_ = 500 nm). Adaptations in this four-opsin visual system are relevant for discrimination of conspecific reflectance spectra in *E. atala*. Together, these findings illustrate how functional changes at multiple rhodopsins contribute to the evolution of a broader spectral sensitivity and adaptation in visual performance.

**Significance Statement:** Rhodopsins are photosensitive protein molecules that absorb specific wavelengths of incoming light and convey colour information in the visual system. We show that molecular evolution in a green insect opsin gene resulted in a shift in its maximal absorbance peak, enabling some lycaenid butterflies to use spectral energy of longer wavelengths (LW) to discriminate colours in the red spectrum better than relatives bearing ancestral green LW rhodopsins. Lycaenids also evolved a duplicate blue opsin gene, and we illustrate an example where species equipped with red LW rhodopsins shifted their blue sensitivity peak to longer wavelengths due to changes in several blue-tuning residues that have evolved repeatedly in different insect lineages. We demonstrate how changes at multiple vision genes in the insect eye effectively create a coordinated mechanism expanding spectral sensitivity for visually guided behaviours such as selecting host plants and mates.

## Introduction

Multiple studies have demonstrated the important contributions of gene duplication and protein-coding changes in the evolution of novelty in lineage-specific phenotypic traits (1–3). For instance, gene duplication combined with modifications leading to functional divergence has contributed to gene family expansion, ultimately increasing transcriptional and functional diversity across lineages (2, 4). Recent examples show that specialization of ancestral functions (5–7) or cooperation between encoded products of duplication (8) are common mechanisms resolving constraints in existing eukaryotic gene networks (9). Genomic segmental duplications have also been shown to promote repeated allelic fixation, linking adaptive duplicated loci to convergent evolutionary pathways (10).

However, structural, functional, or gene network constraints can impose evolutionary trade-offs if there is limited variation for alternative biochemically stable encoded products among duplicated loci within multigene families (9, 11). Characterizing the molecular patterns of evolution within multigene families, and identifying the functional role of mutations at lower taxonomic scales is needed to distinguish the multiple sources of molecular variation. This includes the relative roles of gene duplication, expression changes, protein sequence divergence and convergence, or cellular relocalization of encoded gene products that contribute to new or convergent phenotypes within lineages (2, 7, 8, 12, 13).

Opsins belong to a diverse multigene family of G protein-coupled receptors, offering a robust framework to study how molecular changes can ultimately cause changes in behaviour and favour diversification (14). Opsins bind to a small non-protein retinal moiety derived from vitamin A to form photosensitive rhodopsins and enable vision across animals (15–18). The evolution of vision across the animal kingdom has long been linked to independent opsin gene gains and losses (19–23), genetic variation across opsins (15, 24–26), and spectral tuning mutations within opsins (27, 28). These molecular mechanisms, together with alterations in visual regulatory networks (29), have been shown to contribute to rhodopsin adaptation and the diversification of spectral sensitivity phenotypes in insects (18, 30, 31) and vertebrates including fish, birds, bats and primates (32–36). Thus multiple levels of organization, including the evolutionary diversification of opsin subfamilies, their functional properties and regulatory networks constitute the mechanistic basis for organisms to discriminate light sources of varying wavelengths and ultimately interpret them as colours, yet how multiple changes across opsin paralog repertoires impact spectral sensitivity at small taxonomic scales has been understudied.

The evolution of insect colour vision in particular shows how a complex sensory trait can play a central role in adaptations involving signalling and mate communication (18, 31, 37, 38). Whereas the ancestral repertoire of insects involved three types of light-absorbing rhodopsin genes: ultraviolet (UVRh, 350nm), blue (BRh, 440 nm) and green, also called long-wavelength (LWRh, 530nm) (18), today’s genomes harbor smaller or larger rhodopsin repertoires with strong experimental evidence for repeated functional convergence toward UV, Blue and LW spectral sensitivities across lineages. For example, beetles lost their ancestral blue opsin gene 300 million years ago, and compensated for the loss of blue sensitivity via either UV or LW gene duplication across lineages (23, 39). Blue opsin duplications occurred independently in pierid and lycaenid butterflies (27, 30, 40–42); and extend photosensitivity into the UV/blue in *Heliconius* spp. with λ_max_ = 355 nm and 398 nm (21) and into the violet/blue, in *Pieris rapae* with λ_max_ = 420 and 450 nm (27). UV and LW duplications occurred in butterflies, hemipterans and dragonflies (20, 22, 26, 30, 43–45). In butterflies, LW opsin duplications have been identified in two papilionids, *Papilio xuthus* (31) and *Graphium sarpedon* (46) as well as a riodinid (*Apodemia mormo*) (20, 47), and contribute to extend spectral sensitivity into the far red.

Although duplicated LW opsins have never been detected in the families Nymphalidae, Pieridae or Lycaenidae (25, 40, 41), photoreceptor types with sensitivity peaks in the red have been identified in these groups (40, 41). This supports the hypothesis that additional mechanisms such as lateral filtering and/ or molecular variation of ancestral LW opsin genes also contribute to modify long-wavelength sensitivity.

For example, photostable lateral filtering pigments are relatively widespread across butterfly lineages (e.g. *Heliconius* (21), *Papilio (45), Pieris* (48), *Colias erato* (49), and some moths (*Adoxophyes orana*, (50)*; Paysandisia archon* (51)). These pigments act as long-pass filters, absorbing short wavelengths and pushing the sensitivity peak of LW photoreceptors into the red, to create distinct spectral types that can contribute to colour vision (19, 31, 40, 48, 52, 53). Lateral filtering can shift peak sensitivity to longer wavelengths while reducing peak amplitude, but cannot *extend* photoreceptor sensitivity into the far red beyond the exponentially decaying long-wavelength rhodopsin absorbance spectrum (52). Molecular variation of ancestral LW opsin genes could potentially extend photoreceptor sensitivity, but this mechanism has remained difficult to disentangle from the effects of filtering granules using classical electrophysiological approaches (31, 53, 54).

The molecular underpinnings of mammalian MWS/ LWS spectral tuning has been studied using mutagenesis experiments and led to the identification of critical amino acid replacements and their interactions at key residues (55, 56). Most mammalian lineages possess long-wavelength sensitive (LWS) cone opsins that specify Medium (M, λ_max_ 510-540 nm) and Long (L, λ_max_ > 540 nm) rhodopsins. In humans, trichromatic vision is conferred through the use of Short (S, λ_max_ = 414 nm), and tandem duplicate M (λ_max_ = 530 nm) and L (λ_max_ = 560 nm) cone rhodopsins (34), that collectively allow us to discriminate longer wavelengths of light as green-red colours (57, 58). In birds, tetrachromatic vision is based on two SWS cone opsins, together with a green M opsin (λ_max_ = 497-514 nm) and a red-sensitive LWS opsin (λ_max_ = 543-571 nm) (reviewed in 59). In insects, the evolution of red receptors occurred independently multiple times (30) but the possible contribution of molecular variation to insect LW opsin gene diversification has remained elusive, notably due to difficulties in expressing LW opsins *in vitro* (27, 60, 61, but see 62). Red receptors are intriguingly very common in butterflies compared to other insect groups such as bees or beetles (30), raising the possibility that perception of longer wavelengths plays an important role in the context of foraging (31, 46, 63), oviposition (64, 65) and mate recognition (25) for species equipped with them.

Lycaenids comprise the second largest family of butterflies, representing almost thirty percent of all species, and exhibiting considerable ecological and morphological diversity (66, 67). Pioneering work showed that species of Lycaenidae in the genera *Lycaena* and *Polyommatus* have expanded spectral sensitivity at long wavelengths, and this has been postulated to arise from filtering pigments, modified opsins or both (19, 40, 41). Here, by combining physiological, molecular and functional approaches, we identify additional lycaenid species with red photoreceptors and elevated spectral sensitivity at long wavelengths, and show that their LW opsin locus specifies a novel type of LW rhodopsin with red-shifted maximal absorbance. We focus on the Atala hairstreak (*Eumaeus atala*) as a suitable model to show the interplay between regulatory and adaptive changes at multiple opsin loci in the evolution of red spectral sensitivity and link the evolution of finely tuned four-opsin vision to relevant visually guided behaviours in these butterflies.

## Results

### Hairstreak butterflies with elevated sensitivity at long wavelengths express red-shifted rhodopsin receptors

A photographic series at distinct elevations in the eye revealed dramatic differences in eyeshine colouration between two pairs of lycaenids (Fig. 1 insets and Fig. S1). The Spring Azure, *Celastrina ladon* (subfamily Polyommatinae) and Sheridan’s green hairstreak, *Callophrys sheridanii* (subfamily Theclinae), present homogeneous green and orange eyeshine in their dorsal eyes at 30° elevation, with a relatively small number of red ommatidia (*Ce. ladon*, 5%; *Ca. sheridanii*, 16%) (Fig. S1A-B). By contrast, the Japanese Oakblue, *Arhopala japonica*, and the Atala Hairstreak, *Eumaeus atala* (both subfamily Theclinae), share high levels of saturated red eyeshine in the dorso-equatorial region, due at least in part to species-specific differences in rhodopsin content (Fig. 1, Fig. S1B).

**Figure 1.**
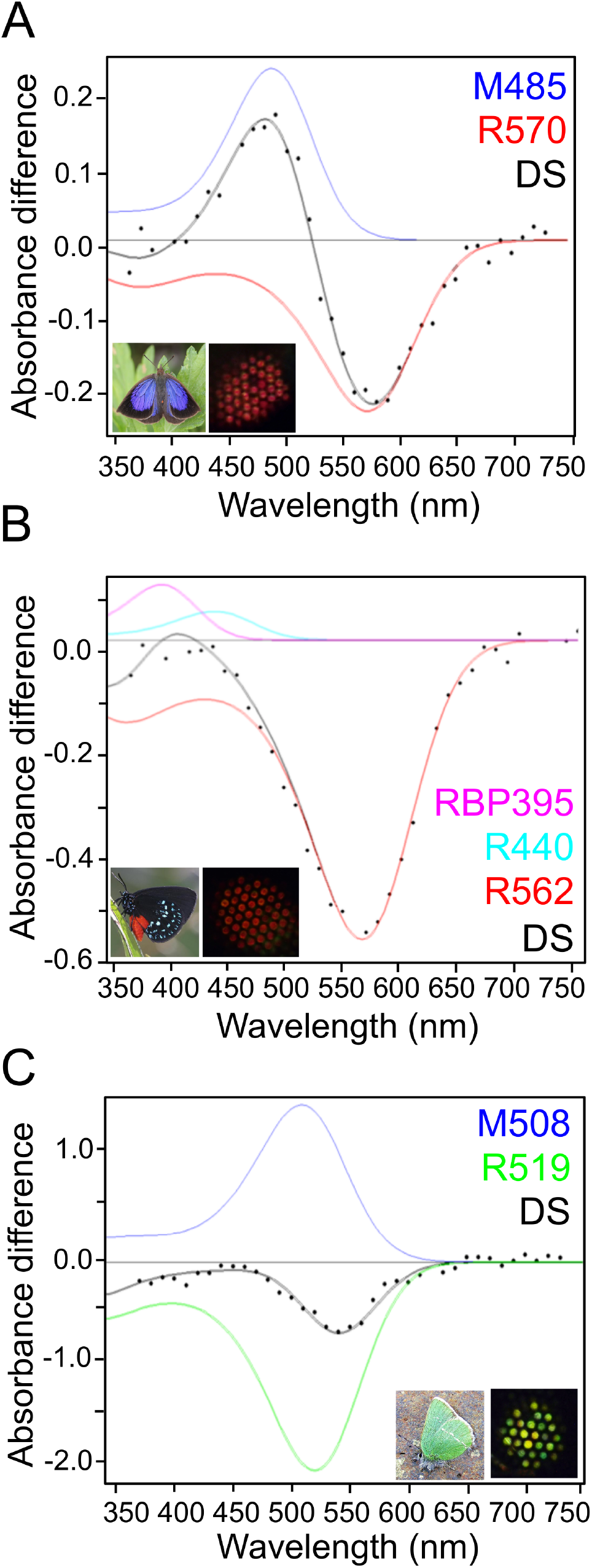
Long-wavelength photochemical difference spectra from lycaenid butterflies. Difference spectra (DS) were obtained following partial bleaches of long wavelength rhodopsins from dorsal retina of intact butterflies. Datapoints represent absorbance differences between amounts of Rhodopsin (R, green or red) and its Metarhodopsin photoproduct (M, blue) product after a dark-period that followed photoconversion. Each black curve represents a computed difference spectrum for least square fits estimates at *(A)* R570 of *Arhopala japonica, (B)* R562 of *Eumaeus atala* and *(C)* R518 of *Callophrys sheridanii*. In *E. atala* the difference spectrum was acquired upon complete degradation of the M photoproduct and with small contributions from R440 and RBP395. In *Celastrina ladon*, λ_max_ of M and λ_max_ of R are very close, which pushed the DS negative peak to the right. Photographs represent respective butterflies (left) and butterfly eyeshine (right) in the dorsal retina (see also Fig. S1).

We partially bleached eyeshine of *A. japonica* and *E. atala* using repeated white-light flashes to reveal two types of ommatidia. Some ommatidia were resistant to bleaching and maintained their red eyeshine owing to lateral filtering by cherry-red pigment granules located distally within the photoreceptor cells of those ommatidia. The rest of the ommatidia were bleached, suggesting the presence of red sensitive opsin-based photoreceptors in these two species (Fig. S1C).

To test for the presence of red sensitive receptors, we performed analyses of *in vivo* photochemical rhodopsin bleaching measurements of adult butterflies. These experiments revealed long wavelength (LW) spectral sensitivities in dark-adapted eyes of *A. japonica* with λ_max_ ± standard error at 571 ± 2.45 nm (CI_95%_ = 566 to 576 nm) (Fig. 1A) and *E. atala* with λ_max_ at 563 nm ± 0.9 nm (CI_95%_ = 561 to 566 nm), respectively (Fig. 1B). We found that the two other lycaenid species we studied, *Ca. sheridanii* and *Ce. ladon*, were difficult subjects for our *in vivo* methods. In *Ca. sheridanii*, a noisy difference spectra obtained by photochemistry provided an estimate at LW λ_max_ = 518 nm ± 3.7 nm for (CI_95%_ = 511 to 526 nm) (Fig. 1C). Similarly, in *Ce. ladon*, partial LW bleaches were not measurable due to low LW rhodopsin densities,.

We chose to examine in detail the *in vivo* and *in vitro* contributions of all rhodopsin pigments in *E. atala*, a multi-brooded and abundant hairstreak butterfly naturally occurring throughout the year in Florida (USA) (68). Unlike most lycaenids, *E. atala* larvae are extreme specialist herbivores on New World cycads in the genus *Zamia* and show an unusually bright aposematic colouration advertising toxins they sequester from their hosts (69). Conversely, the adult is a striking velvety black butterfly with bright blue and red colours on the wings and the abdomen (Fig. 1) and can collect nectar from 43 species across 20 plant families (69)

We first analyzed epi-microspectrophotometric difference spectra obtained after intense ommatidial flashing from a series of interference filters, which in addition to identifying R565 (LW rhodopsin, Fig. 1B) allowed us to narrow down a blue rhodopsin at R440 (Fig. 2A). Optophysiology measurements confirmed that neither the R565 LW nor the R440 blue sensitive rhodopsins could be responsible for the high sensitivity around 360 nm, which instead was due to UV-receptors (Fig. 2B, Table S2). Finally, a densitometric analysis of male and female eyes (Fig 2C-D, Tables S3-S4), showed that these physiological data are best fit using least-squares regression by a model in which four rhodopsins are present in the eye with λ_max_ values matching the LWRh rhodopsin as well as UVRh 360 nm, BRh1 441.3 nm ± 4.7 nm (CI_95%_ = 431.7 to 450.9 nm, Table S2) and a second blue rhodopsin BRh2 494.2 ± 1.2 nm (CI_95%_ = 492 to 497 nm) (Fig. 2E-F, Tables S3-4).

**Figure 2.**
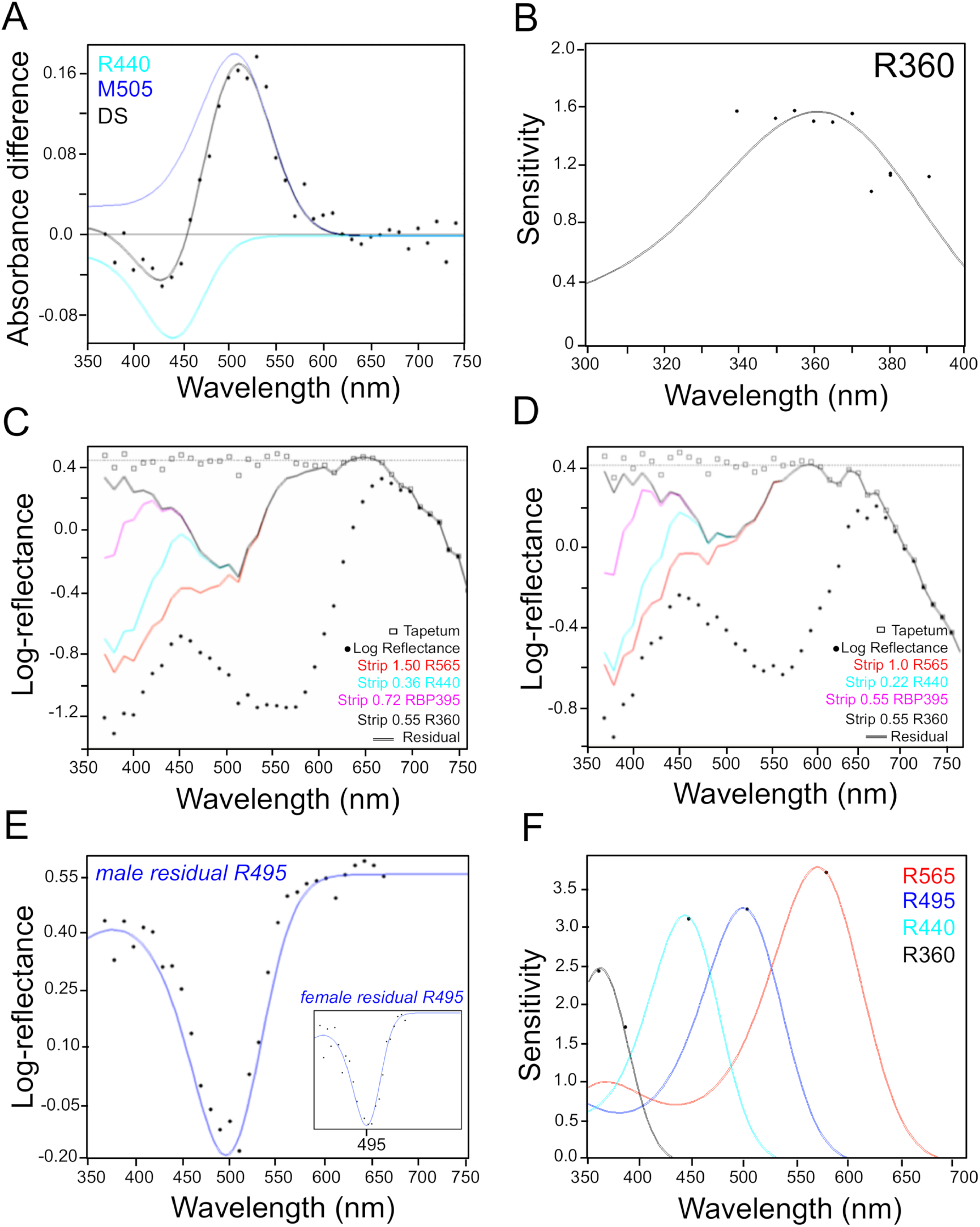
*In vivo* evidence for four rhodopsins in the Atala hairstreak butterfly. *(A)* Photochemical analysis of a butterfly male eyeshine using an epi-microspectrophotometer. Log-reflectance difference absorbance spectra (DS, Difference Spectrum, filled circles) and fitted curves (solid line) measured from dark-adapted eyes via partial-bleaching experiments of R440. *(B)* Optophysiological analyses designed to measure pupillary sensitivity in the UV. The least squares fit analysis provides confident estimates for a UV rhodopsin with λ_max_ at 360nm. Comparison to sensitivities in the blue, driven by R440 and in the red, driven by LW R565 rhodopsins, show that those two opsins make no contributions in the UV. *(C-D)* Densitometric analysis of an epi-microspectrophotometric reflectance spectrum of a male *(C)* and female *(D)*. Completely dark-adapted *E. atala* eyeshine is used to confirm the contribution of each estimated visual pigment in the eye. The black dots plot the log-reflectance spectrum. The red curve is the spectrum after having computationally stripped optical density (OD) 1.50 of rhodopsin with λ_max_ 565 nm so that the residual spectrum is flat from 565 nm to 660 nm; cyan curve, the log-reflectance spectrum after having stripped OD 0.36 of R440 to produce a residual flat from 450 nm. The black curve is the log-reflectance spectrum after stripping OD 0.72 for R395 and 0.55 R360, leaving a residual spectrum fit by a fourth-rhodopsin R495. The densitometric analysis of female eyes *(D)* is qualitatively identical, with stripping densities of 1.55 R565, 0.40 R440, 0.80 RBP395 and 0.55 R350, leaving a residual indicative of a fourth pigment contribution, fit by rhodopsin R495. Stripping the R495 residual leaves the curve plotted in open squares that is the putative average log-reflectance spectrum of the tapetum, that is flat from the UV out to 660 nm. *(E)* Computational analysis of male residual reflectance spectra supports the contribution of a fourth opsin pigment with λ_max_ peaking around 495 nm. The residual fit for females is shown in the inset. *(F)* Sensitivity data from an optophysiological threshold experiment measured at wavelengths close to λ_max_ values (360 nm, 440 nm, 495 nm, 562 nm) of the four rhodopsins. This shows that sensitivities at 440 nm and 495 nm cannot possibly be driven by R565.

### Red sensitivity is due to functional variation at long-wavelength opsin loci

Transcriptomic mRNA profiling of *E. atala* yielded only a single LW opsin (Table S6), which is in line with earlier molecular evidence showing that lycaenid species possess one LW opsin gene (30, 40, 53). Accordingly, eye cDNA library screening using degenerated oligonucleotide primers followed by RACE cDNA amplification led to the characterization of single orthologous LW cDNA sequences in *Ce. ladon, Ca. sheridanii* and *A. japonica* (Fig. 3). We optimized an *in vitro* HEKT293 cell culture assay to reconstitute heterologous rhodopsin pigments (27, 61, 70, 71) by using a newly engineered heterologous vector to increase expression levels, and improving purification procedures to obtain higher yields of actively reconstituted LW rhodopsins. Our expression cassette derives from the pcDNA5 plasmid and expresses coding opsin sequences tagged with a C-terminal FLAG epitope under a strong CMV promoter. Directly downstream the FLAG epitope, a short peptide linker plus a T2A cleavage cassette enable co-transcription of a fused cytoplasmic fluorescent mRuby2 coding sequence (Fig. 3A). This expression cassette proved to be efficient at detecting monomeric units of ultraviolet, blue and long-wavelength (LW) heterologous *E. atala* opsin proteins *in vitro* (Fig. S2A). It is also successful at reconstituting and purifying active LW rhodopsins, *i.e*. in *A. japonica, E. atala, Ca. sheridanii* and *Ce. ladon* lycaenid butterflies (Fig. 3B-F, Fig. S2B, Table S5).

**Figure 3.**
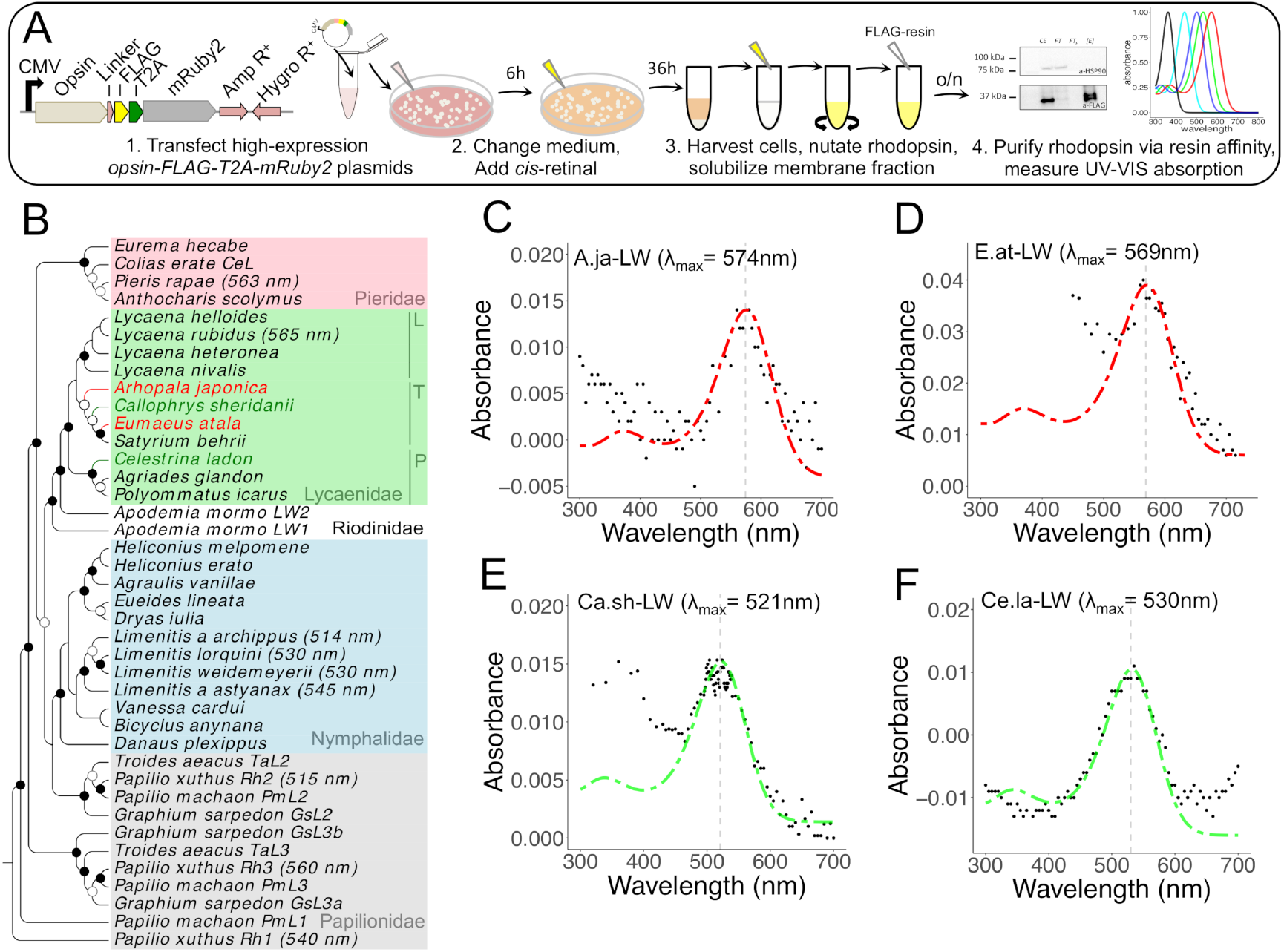
Functional characterization of red-shifted long wavelength lycaenid butterfly opsins. *(A)* Schematic of modified pCDNA expression vector cassette and workflow for functional characterization of rhodopsin complexes with cis-retinal chromophore. All steps following addition of cis-retinal are performed under dim red-light illumination. *(B)* Neighbour-Joining (NJ) tree of butterfly long wavelength opsins. Bootstrap node support is as follows: 50-74%, white circle; 75-94%, grey circle; ≥ 95%, black circle. Previously published λ_max_ physiological data are indicated in parentheses when known (20, 27, 40, 41, 45). Corresponding family names are labeled on the right. L, T and P correspond to lycaenid subfamilies Lycaeninae, Theclinae and Polyommatinae, respectively. *(C-F)* Dark spectra of long wavelength rhodopsin (LWRh) expressed using an HEKT293 transient cell culture system and purified via FLAG-epitope. LWRh rhodopsin absorbance spectra are indicated with black dots (Table S7, Dataset S1), and a rhodopsin template data (100) was computed to obtain the best estimates of λ_max_ fitting the data. (C) *Arhopala japonica* purified LW opsin with λ_max_ = 570 nm, (D) *Eumaeus atala* purified LW opsin with λ_max_ = 567 nm, *(E) Callophrys sheridanii* purified LW opsin with λ_max_ = 525 nm, (F) *Celastrina ladon* purified LW opsin with λ_max_ = 525 nm.

When purified from large-scale HEK293T cell cultures and reconstituted *in vitro* in the dark in the presence of 11-*cis*-retinal, we found that the LW rhodopsin from *A. japonica* absorbs maximally at λ_max_= 574 ± 4 nm (CI_95%_ = 570-586 nm) (Fig. 3C) whereas that of *E. atala* absorbs maximally at λ_max_= 569 ± 2 nm (CI_95%_ = 565-573 nm) (Fig. 3D). The peak of absorbance of purified LW rhodopsin measurements is within confidence intervals of the best fit for LW linear absorbance estimates *in vivo* and supports the hypothesis that the LW rhodopsin limb of absorbance in these two species pushes red sensitivity above 600 nm. The LW rhodopsin pigment from *Ca. sheridanii* absorbs maximally at λ_max_ = 519.2 nm ± 1.1 (CI_95%_ = 517-521 nm) (Fig. 3E), which is in accordance with photochemical measurements (Fig. 1C), whereas *Ce. ladon* LW rhodopsin absorbs maximally at λ_max_= 531.7 ± 1.5 nm (CI_95%_ = 529-535 nm) (Fig. 3F, Table S5).

In summary, these findings indicate that our expression system can be used successfully to assess the functionality of LW rhodopsins outside of the complex eye environment and demonstrate that at least some hairstreaks (Theclinae) express a new functional type of far-red shifted visual opsin.

### Blue opsin mRNA expression patterns lead to enhanced eye spectral richness

Following photochemical and densitometric evidence that *E. atala* butterflies likely possess four rhodopsins including contributions from two blue rhodopsins, we identified two differentially expressed blue opsin-like transcripts from the eye transcriptome, namely BRh1 and BRh2 (Table S6). We then examined their respective mRNA expression patterns in photoreceptor cells, aided by the histological reconstruction of photoreceptor organization in a typical ommatidium (Fig. 4).

**Figure 4.**
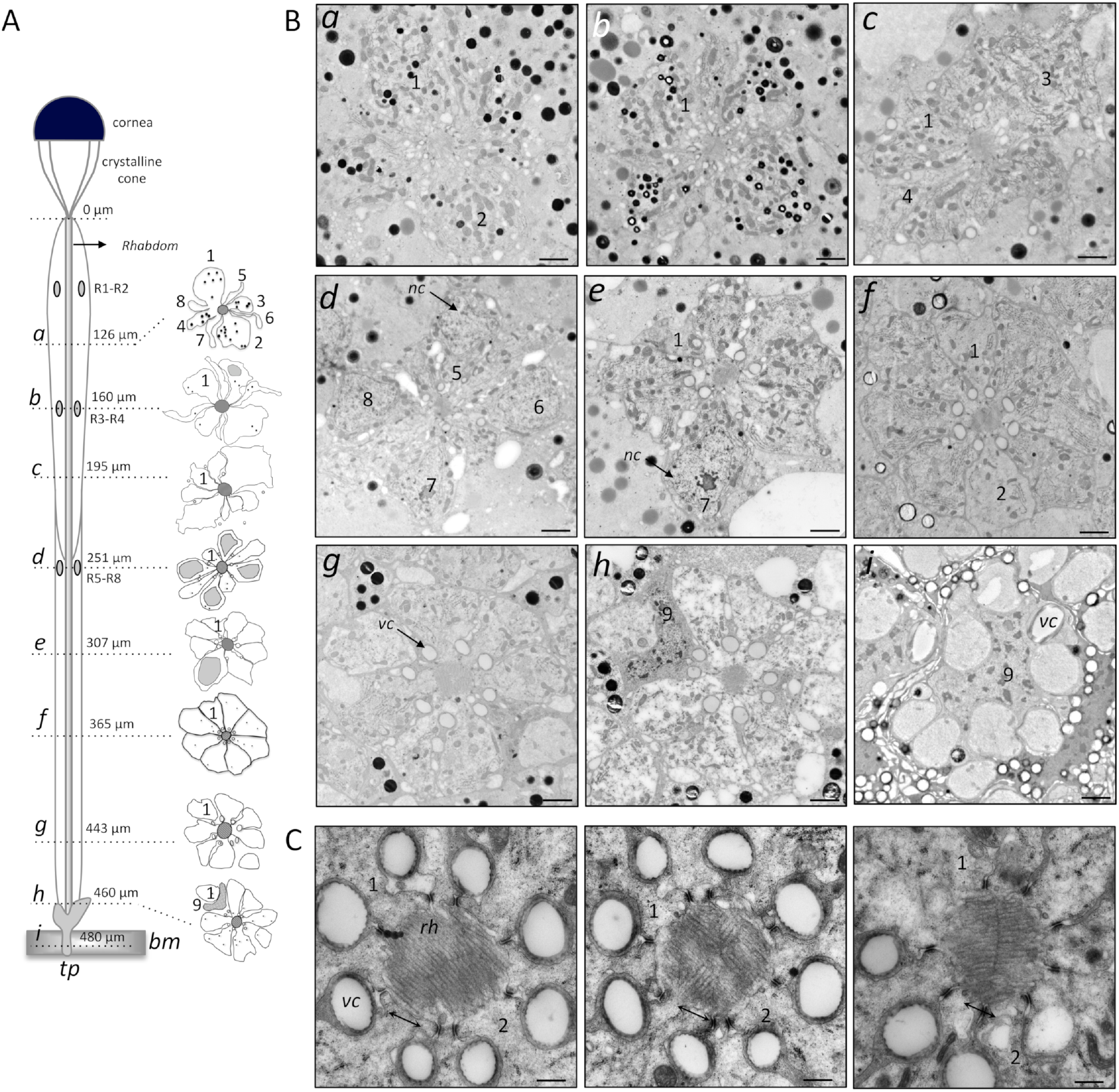
Anatomical overview of a typical ommatidium in *Eumaeus atala*. *(A)* Diagram illustrating a typical ommatidium and the relative contribution of nine photoreceptor cells R1-R9 at different depths along the rhabdom (480 μm long). Photoreceptors R1 and R2 are proximal cells in region *a* (126 μm) that contribute microvillar structures containing UVRh, BRh1 and BRh2 rhodopsins. Photoreceptors R3-R4 are proximal receptors containing LWRh and Brh1 rhodopsins. Photoreceptors R5-R8 are distal cells exclusively expressing the LWRh rhodopsin (see also Fig 4, Fig S2). The basal cell, R9 is restricted to the region immediately proximal to the basement membrane (460 μm, h). The expressed mRNA opsin type was not investigated in this cell. *(B)* Scanning electron micrographs from a male dorsal eye region at 30° elevation across the rhabdom. Scale bars, 2 μm. *(C)* Distal microvilli in regions *a-b* are exclusively oriented parallel to the R1-R2 axis (left panel). As R3-R4 cells expand and contribute the majority of microvilli from regions *c* to *e*, together with novel contributions from proximal receptors R5 to R8, three ommatidia subtypes are formed that exhibit microvillar contributions parallel to (left panel) but also intertwisted (middle panel) and perpendicular to the R1-R2 axis (right panel). Scale bars, 500 nm*. bm*, basement membrane*; rh*, rhabdom*; nc*, nucleus*; tp*, tapetum*; vc*, vacuole; 1-9, photoreceptor cells R1 to R9.

We found that *E. atala* hairstreaks have a straight, 480-micron long rhabdom composed of eight longitudinal photoreceptor cells (Fig. 4A-B (a-g)) and a ninth cell close to the basement membrane (Fig. 4A-B (h)). The two most distal R1-R2 photoreceptor cells contribute the majority of microvillar extensions from 0 to 160 microns, whereas R3-R4 distal cells contribute a majority of microvilli from 140 to 300 microns, thereby overlapping partially with R1-R2 in the distal rhabdom tier. The proximal R5-R8 cells contribute most microvilli in the last rhabdom tier up to 440 microns, a depth where the photoreceptor cells no longer bear any microvilli and the ninth cell becomes visible (Fig. 4A-B (i)).

This rhabdomeric analysis provided the spatial morphological insights necessary to subsequently identify photoreceptor cells expressing each opsin mRNA across different eye regions. Using double fluorescent *in situ* hybridization experiments in transverse and longitudinal eye sections of males and females of *E. atala*, we showed that LWRh is expressed in all ommatidia in the six photoreceptor cells R3 to R8 (Fig. 5), which is typical of many butterfly species (25, 29). No fluorescent signal was detected for the long-wavelength opsin in R1-R2 cells (Fig. 5D-F). We next examined the cellular localization of the short UV and medium blue opsin mRNAs in transverse sections in the dorsal eye. Using probes targeting LWRh in combination with cRNA probes for UVRh, BRh1 or BRh2 mRNAs, our data provide evidence that the latter rhodopsin mRNAs can be expressed in either or both R1-R2 receptor cells forming single ommatidia (Fig 5A-F).

**Figure 5.**
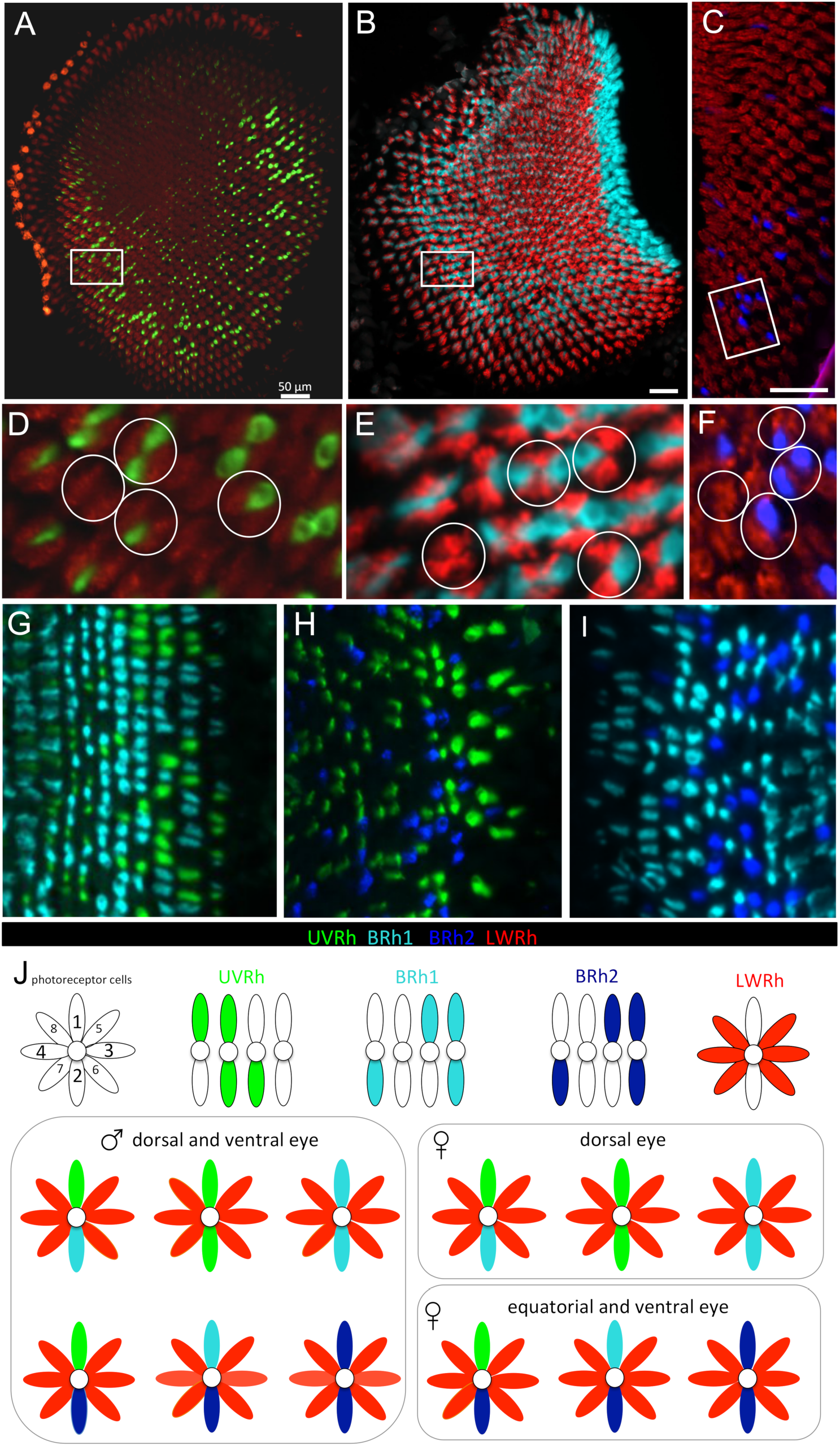
Adjacent photoreceptor localization between duplicate blue opsin mRNAs drives retinal mosaic expansion. Double fluorescent *in situ* hybridization shows six ommatidial types in *E. atala* compared to ancestral butterfly eyes. *(A-F)* Exclusive one-cell one-mRNA expression pattern in distal photoreceptors R1 and R2, showing for *(A)* UVRh (inset in *(D), (B)* BRh1 mRNA (inset in *(E)*) and *(C)* BRh2 mRNA (inset in *(F)*). Circles highlight the four distinct opsin expression patterns in R1 and R2. *(G-I)* UVRh, BRh1 and BRh2 mRNAs do not coexpress in R1-R2 cells. Photoreceptor cells R3-R8 express the LWRh opsin (red). Males and females show dorso-ventral dimorphism (Fig. S4). *(J)* Schematic representation of opsin-based photoreceptor classes in males and females of the six ommatidial types as follows (R1-R2-R3/R8): UV-UV-LW, UV-B1-LW, UV-B2-LW, B1-B1-LW, B1-B2-LW, B2-B2-LW.

We also assessed the possibility that these opsins are co-expressed in R1 and R2 cells using cRNA probes for UVRh and BRh1 (Fig. 5G), UVRh and BRh2 (Fig. 5H) or BRh1 and BRh2 (Fig. 5I). We find that R1 and R2 follow a one-cell one-opsin regulation pattern, with mutually exclusive expression of UVRh, BRh1 and BRh2 mRNAs (Fig. 5J). Females exhibit a dorso-ventral expression gradient in which BRh2-expressing cells are sparse in the dorsal eye region. In the ventral retina however, all three opsins (UVRh, BRh1, BRh2) are found in addition to LWRh (R3-R8) (Fig. S3).

Cellular expression data therefore show that both of the duplicated blue opsin mRNAs encode functional transcripts. Overall, the cellular localization of blue opsin duplicates in adjacent photoreceptor cells creates multiple ways in which rhodopsins are distributed within individual ommatidia, forming a local stochastic rhodopsin mosaic of at least six opsin-based photoreceptor classes (UV-UV, B1-B1, B2-B2, UV-B1, UV-B2, B1-B2, Fig. 5j). Additional structural features of the eye may also contribute to the diversity of spectral sensitivity functions of individual photoreceptors, including the spectral influence from lateral filtering granules found in distal rhabdomeres that add to the spectral sensitivity of some photoreceptor cells.

### BRh1 and Brh2 opsin duplicate loci encode blue and green-shifted rhodopsins

We reconstituted active BRh1 and BRh2 rhodopsins *in vitro* and measured their spectral sensitivities via spectroscopy as detailed in the methods. We determined that BRh1 λ_max_ = 435 nm ± 2 nm and BRh2 λ_max_ = 500 nm ± 2 nm (Fig. 6A-B, Table S7), thereby confirming that they encode the rhodopsins conferring the expanded blue to green spectral sensitivity in *E. atala*. In addition to its LWRh red-shifted opsin (Fig. 3D), *E. atala* uses a fourth rhodopsin that absorbs UV wavelengths *in vitro* at λ_max_= 352 nm ± 3.5 nm (CI_95%_ = 345-360 nm) (Fig. 3, Fig. S2d, Table S7).

**Figure 6.**
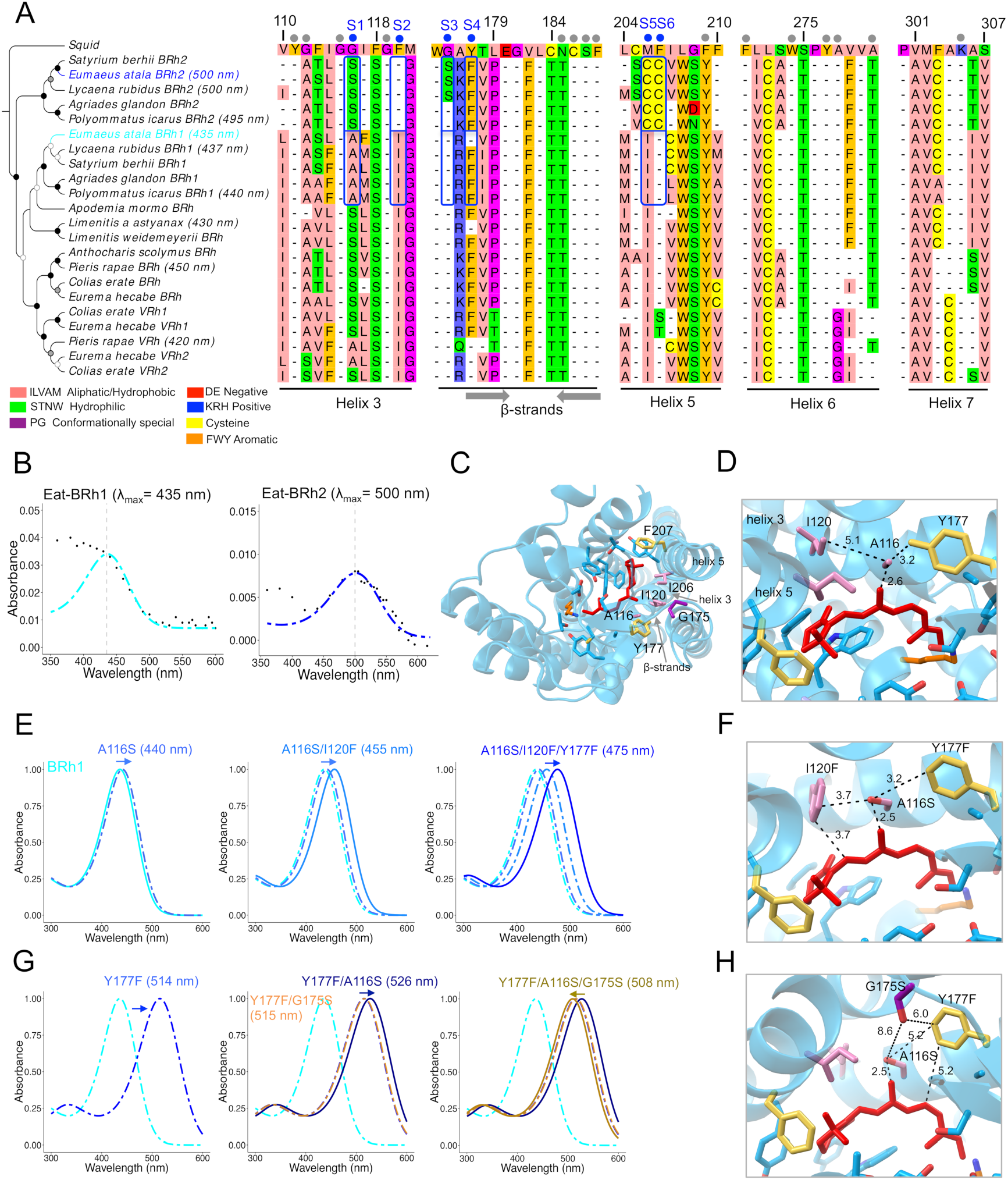
Residues responsible for spectral tuning shifts in blue rhodopsins. *(A)* NJ tree of selected lepidopteran blue opsin amino acid sequences. The squid rhodopsin, *Todarodes pacificus* (acc. nr. CAA49906) is used as outgroup. Bootstrap node support is as follows: 50-74%, white circle; 75-94%, grey circle; ≥ 95%, black circle. Dots above the partial multiple sequence alignment shows the 21 amino acid residues residing within 5 Å of any carbon atom in the retinal polyene chain. *Blue* dots identify the six positions where amino acid residues differ between *E. atala* BRh1 and BRh2, blue rectangles highlight variants at these positions in lycaenids. Residue numbering is based on residue position in the squid rhodopsin. Residues are coloured according to their physicochemical properties in Jalview v2 (105). Grey arrows indicate β-strands forming the binding pocket. *(B)* Blue opsin absorbance spectra (dots) fitted to the visual template (cyan and blue line functions), respectively. *(C)* Predicted structure for Eat-BRh1 based on homology modeling with the squid rhodopsin with variant sites Ala116, Iso120, GLy175, Phe177, Iso206. *(D)* Native Brh1 rhodopsin bearing S1 (A116), S2 (I120) and S4 (Y177). *(E)* Substituting residues 116, 120 and 177 leads to a 30-nm partial blue-shift in rhodopsin absorbance λ_max_ in the triple mutant (see also Figure S2). This tuning shift may be mediated by several possible mechanisms including additive effects caused by novel hydrogen bond formation at the coevolving adjacent sites 116 and 120 *(F)* and with nearby conserved residues G115 and G121. *(G)* An alternative evolutionary route involves substituting G175S (S3) which partially compensates the green tuning shift of double mutant A116S/Y177F (Fig. S6, Table S8) and tunes the absorbance spectrum near 500nm. These alternative trajectories highlight additive and epistatic interactions between four residues at sites 1-4 *(F, H)* in the acquisition of the blue-shifted function.

### Four spectral tuning sites and epistasis leads to bathochromic shifts between blue duplicates

In order to understand the proximate mechanisms driving the 65 nm spectral shift between blue opsin duplicates, we performed homology modeling against the invertebrate squid rhodopsin (72). Along with their 380-aa long opsin protein sequences, Eat-BRh1 and Eat-BRh2 exhibit 101 amino acid residue differences (corresponding to 73% similarity in aa sequence). Six variant sites were identified among 21 homologous residues located within 5Å of any carbon forming the *cis*-retinal binding pocket (Fig. 6A,C). Two variant sites (S1 and S4) are shared with a blue/violet opsin duplication that occurred independently in pierid butterflies, causing a UV-shift (VRh λ_max_ = 420 nm) from the ancestral blue rhodopsin (BRh λ_max_ = 450 nm) (27), whereas four residues are unique to duplicated blue opsins found in lycaenids (S2, S3, S5,S6).

Given this combination of shared and unique yet limited number of variant sites, we decided to test their possible involvement in blue spectral tuning by site-targeted mutagenesis. Specifically, we first modified individual residues S1 to S4 and measured their spectral tuning effect *in vitro* (Fig. 6E-G, Fig. S4). Chimeric opsins showing bathochromic shifts were used sequentially to substitute adjacent variant sites following two candidate evolutionary trajectories. We observed an additive bathochromic shift totaling 40 nm (λ_max_ = 475 nm) by substituting A116S (S1) together with I120F (S2) and Y177F (S4) (Fig. 6E). In a second tuning trajectory, we observed that Y177F alone conferred a 79 nm bathochromic shift (λ_max_ = 514 nm), which could then be compensated by a 6 nm hypsochromic shift (λ_max_ = 508 nm) by combining G175S and A116S (Fig. 6G). A third evolutionary trajectory explored the contribution of two lycaenid-specific cysteine substitutions (I106C, F207C) in helix 5 (Fig. S4). These changes caused a strong green-wave spectral shift that was not compensated for by additional candidate interacting residues in the quintuple mutant (S1,2,4,5,6; Fig. S5, Table S8). G165S (S3) tested alone or in various double mutant combinations caused bathochromic shifts, but we did not obtain the sextuple chimeric construct bearing G175S (S3) and cannot therefore exclude the possibility that it also plays a role in this case. The available results from all variants bearing cysteines in S5 and S6, however, suggest that both cysteine residues on helix 5, which are absent in Pieridae blue opsins, are unlikely to be the main evolutionary drivers of blue spectral shifts in lycaenid BRh2 loci.

Finally, our optophysiological density analyses showed that blue spectral sensitivity differs between species equipped with green or red LW rhodopsins. Whereas all species possess a blue photoreceptor with conserved absorbance with λ_max_ = 435-440 nm, the second blue sensitivity peak in A. japonica and E. atala is at λ_max_ = 500 nm (Fig. 2F, S5A) unlike λ_max_ = 489 in C. ladon (Fig. S6B, Table S9). These data suggest a role for coordinated shifts between blue and LW rhodopsins in the evolution of red spectral sensitivity.

### Shifted Blue and LW rhodopsins tune colour vision in the context of conspecific recognition

In order to investigate possible correspondence between *E. atala* visual spectral sensitivities and colour traits that might be important in signalling and sexual selection, we measured the reflectance spectra associated with specific wing patches (Fig. 7, SI Dataset 2). For the butterfly to interpret colours, it must i) possess at least two spectral types of receptors sensitive to the reflectance spectrum of incident visible light illuminating the coloured area, and ii) be able to compare individual receptor responses neurally to create an output chromatic signal (73). Although we did not take this analysis to the second step of recording responses of different individual receptors to reflectance spectra, we investigated in detail the first necessary condition by measuring and comparing reflectance spectra of body and wing patches from males and females including blue scales on the abdomen and thorax, as well as black, blue and red scales on forewings and hindwings (Fig. 7A). In males, dorsal forewings are bright iridescent blue in summer, whereas scales appear more generally green/ teal in winter generations (68). Female dorsal wings, on the other hand, display a darker royal blue colour along the edge of their upper forewings. Both sexes also have conserved wing and body patterns, including regularly spaced rows of blue spots visible on folded and unfolded hindwings, and a bright red abdomen with a large bright red spot on the mid-caudal hindwing area that falls precisely along the abdomen in folded wings (Fig. 1A, inset).

**Figure 7.**
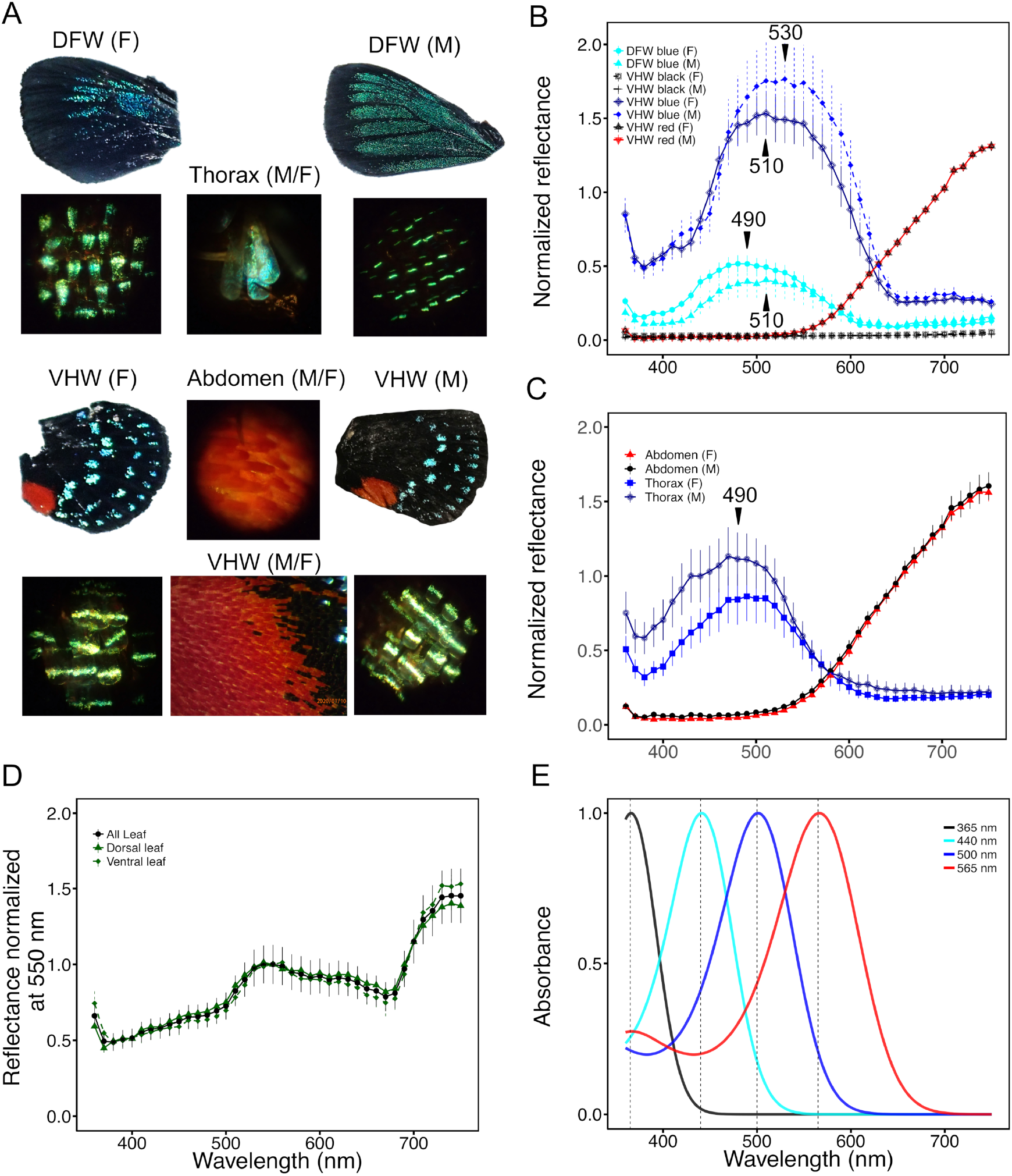
Wing and body spectral reflectance in *Eumaeus atala*. *(A)* Female (F) and male (M) dorsal forewing (DFW), ventral hindwing (VHW), thorax and abdominal scales. The photographs below each wing are magnified views of representative coloured wing scales in patches measured by epi-microspectrophotometry (MSP). The field of view for each photograph is 210 μm in diameter. Photographs were acquired using an Olympus TG-1 camera set at zoom 2.8. *(B-C)* Graphs of mean reflectance spectra ± standard errors of the mean of wing scale patches *(B)* and body scales *(C)* (N=3-5 individuals per sex). *(D)* Leaf reflectance spectra (4-5 measurements per leaf surface) (SI Datafile 2). *(E) E. atala* rhodopsin absorbance spectra. Male and female wing reflectance spectra are not exclusive, and spectral shapes are highly consistent indicating that male and female spectra are similar in the visible. In females, hindwing cyan spots show a reflectance peak at 540 nm *(B)*. Blue scales on the thorax and dorsal forewings have a reflectance peak at 490-500 nm. Hindwing *(B)* and abdominal red patches *(C)* have identical high reflectance spectra at long wavelengths in both sexes. MSP analyses from black areas of ventral hindwings *(B)* showed that black scales are only 1 % as reflective as adjacent cyan scales in the blue/green band. Brightness in black regions decreases 100-fold compared to adjacent cyan scales in both sexes. All reflectance curves overlap at contributing wavelengths for at least two rhodopsins, thereby efficiently distinguishing foliage and sex-specific wing colour patterns.

First, epi-microspectrophotometry measurements from dark areas of male and female wings (Fig. 7B) showed that black scales are only 1 % as reflective as adjacent cyan scales in the blue/green band at 450 nm, with a reflectance maximum of 0.053 (female, *f*) and 0.041 (male, *m*) (SI datafile 2) compared to reflectance maxima at 510 nm of ventral hindwing blue scales 1.53 (*f*) and 1.76 (*m*) (Fig. 7B). This indicates that brightness – which is a function of reflectance - in black regions decreases 100 fold compared to adjacent coloured scales in both sexes.

Blue scales on the dorsal forewings have a maximal reflectance peak at 490 nm in females and 510 nm in males, and blue scales on the ventral hindwings have a maximal reflectance peak at 510 nm (*f*) and 530 nm (*m*) (Fig. 7B). Thorax scale reflectance is 1.86 times and 2.88 times higher than dorsal wing reflectance in females and males, respectively.

Red scales on male and female hindwings reflect maximally in the far red (750 nm), similarly to abdominal scales (Fig. 7B-C). Blue scales on the thorax have a maximal reflectance peak at 490 nm in both sexes, overall indicating that male and female reflectance spectra are similar for blue scales on the body, but not for blue scales on the wings. We noted, too, that leaf surfaces of the butterfly’s primary host cycad, *Zamia integrifolia*, have a peak of reflectance at 550 nm and a red edge inflection point around 700 nm with high reflectance in the near infrared region (Fig. 7D). Altogether, our analyses support that the butterfly’s photoreceptor spectral sensitivities (Fig. 7E) can efficiently discriminate between its host plant (for oviposition), the colours of male and female conspecifics, and colour variation between sexes.

## Discussion

### Red opsin receptors contribute to far-red spectral sensitivity

It has remained challenging to identify the contribution of different genetic mechanisms that affect phenotypic variation amongst complex traits in nature, and ultimately an organism’s fitness. Opsins represent a robust system to link molecular changes to phenotypic changes in animal colour vision as they are the first elements in phototransduction cascade and have been shown to directly modulate the visual spectral sensitivity of insects, primates and other vertebrates.. We have optimized a functional assay to disentangle the contribution of the long-wavelength (LW) rhodopsin from filtering pigment granules and variable eye reflectance properties in the eyes of lycaenid butterflies. We discovered a novel functional form of LW rhodopsin with a red-shifted maximal absorbance spectrum (λ_max_) between 565 and 575 nm in two lycaenid species, *A. japonica* and *E. atala*, that is further supported by microspectrophotometry (MSP) and optophysiology data (Figs. 1–3, Fig S5). The red-shifted rhodopsins have a longer-wavelength limb of absorbance than lycaenid green LW rhodopsins due to a bathochromic shift in their λ_max_ (Figs. 1, 3), which elevates the photoreceptor response at long wavelengths (Fig. 2F, S5A). Consequently, those two species have much greater optophysiological sensitivity in the far red than the lycaenid species with photoreceptors expressing green LW rhodopsins, whether or not surrounded by lateral filtering pigments.

Red spectral sensitivity has previously been examined *in vivo* notably in a species of cabbage white butterfly, *Pieris rapae*, where light perception above 600 nm results from ommatidia expressing a single LW opsin together with two types of pigment granules that confer three types of red photoreceptors (48). Methods employing cAMP-dependent heterologous spectroscopy identified maximal peak absorptions at 540 nm and 560nm for *Papilio* PxRh1 and PxRh3 LW opsins respectively (62), whereas earlier electrophysiological λ_max_ estimates placed PxRh1 at 525 nm and PxRh3 at 575 nm (45). Our assay provides quantifiable expression and yields to obtain accurate λ_max_ *in vitro* outside of the complexity of the eye structure itself which circumvents potential inaccuracy from optophysiological estimates measured *in vivo* in certain species due to interfering lateral pigments or other properties of the eye itself. It also has the advantage of functionally studying variation in orthologous rhodopsin genes independently from inferences based on sequence data alone. Our study reconciles *in vivo* and *in vitro* approaches to substantiate that higher visual performance in the far red is achieved by modifying absorbance properties of long-wavelength rhodopsins.

### Duplication, spectral tuning and adjacent cellular localization provide expanded photoreceptor types

Opsin evolution has undergone recurrent events of gene duplication and loss across animals including insects (16). From ancestral insect trichromatic colour vision (UV, blue, green), insects that have lost one of these three rhodopsins are often found to have compensated from that loss by recruiting a duplicate gene copy that has undergone spectral tuning to shift the peak sensitivity, as seen in true bugs and beetles (23, 39). By contrast, tetrachromatic insects that activate an additional rhodopsin are also known to have acquired increased spectral sensitivity in ranges of visible light, as seen in *Heliconius* duplicated UV opsins (21), or *Pieris* duplicated blue opsins (27). Lycaenid species examined thus far have two blue opsins for which epi-microspectrophotometric estimates previously yielded sensitivities of λ_max_ in the range 420-440 nm and 480-500 nm (40, 41).

Our results show that blue opsin loci in *E. atala* specify blue and green-shifted rhodopsins (Figs 2,6) similarly to species of *Lycaena* and *Polyommatus* (40, 41). The most striking functional insight in the evolution of blue spectral tuning in *E. atala* comes from chimeric BRh1 variants bearing mutations A116S, G175S and Y177F, which confer a 73-nm bathochromic shift (λ_max_ = 508 nm) that most closely recapitulates the spectral properties of Eat-BRh2 (λ_max_ = 500 nm) compared to other tested variants (Fig. 5G, Fig S4). Intermediate adaptive phenotypes can also be revealed via gradual evolutionary trajectories. Eat-BRh1 variant A116S causes a +5 nm shift alone, but together with I120F, shifts maximal absorbance by an additional +15 nm to λ_max_ = 455 nm. Since these two residue substitutions are conserved across all characterized lycaenid BRh1 loci (Fig. 3A), these results show how adjacent sites in helix 3 can contribute to intermediate blue absorbance spectra across lycaenids.

The third tuning residue, Tyr177Phe is a key spectral tuning mutation in *E. atala*, since the triple BRh1 variant (A116S/I120F/Y177F) displays a 30 nm bathochromic shift (λ_max_ = 475 nm) compared to its native rhodopsin, and illustrates the multiple ways that gradual spectral tuning can evolve, at least in this species. Two of the reverse tuning substitutions are the same sites responsible for hypsochromic spectral shifts both in a blue-shifted LW rhodopsin of a *Limenitis* butterfly (Y177F, −5 nm) (61) and in a violet-shifted blue rhodopsin of a *Pieris* butterfly (S116A, −13 nm; F177Y −4 nm) (27), stressing the importance of tuning residues lying on the ionone ring portion of the chromophore binding pocket. Spectral tuning modulation in blue-rhodopsin duplicate functions has therefore involved conserved biochemical constraints along independent evolutionary trajectories that selected for partial spectral tuning sites in rhodopsin G-coupled receptors. Reverse mutations are not functionally equivalent in their absolute magnitudes (Δλ_max_), therefore underscoring the role of epistatic interactions with neighbouring sites resulting in distinct λ_max_ shifts across butterfly lineages.

Molecular and phylogenetic studies of opsin evolution in vertebrates have also shown that homologous tyrosine residues in the ionone ring portion of the chromophore-binding pocket (e.g. Y262) in the human blue cone opsin (λ_max_ = 414 nm) are responsible for a 10 nm bathochromic spectral tuning when mutated to Tryptophan (Trp) (74). This illustrates that distant opsin loci meet similar structural and biochemical constraints as those observed in the evolution of vision genes in other vertebrates (11, 14). However, Y177 is unique to *E. atala*, and other BRh1 loci at this position readily possess the F177 similarly to Eat-BRh2 (Fig. 3A). Not all BRh2 loci have S175 but instead keep G175 in both blue loci, suggesting an additional yet unknown role for adjacent residues in positions R176K and I178V. The latter residues do not differ highly in hydrophobicity but may still provide the necessary molecular interactions with the ionine ring portion of the chromophore necessary to modulate variable BRh2 spectral phenotypes. Human ancestors also achieved blue sensitivity gradually and almost exclusively via epistasis at seven amino acid residues (55). By studying independent opsin gene duplicates in the butterfly family Lycaenidae, our *in vitro* assays refine our understanding of the molecular basis of convergent colour vision phenotypes and help to identify key determinants of genotype-phenotype relationships across insect blue opsins, although these are only a snapshot of all possible chimeric blue rhodopsin variants along at least three possible evolutionary trajectories. Testing the non-additive interactions at co-evolving BRh2 adjacent sites (175-178) in lycaenids lacking red sensitivity will help to fully recapitulate intermediate phenotypes across derived blue-shifted rhodopsin duplicates in addition to those generated by A116S and I120F.

In theory, a typical tetrachromat can achieve better wavelength discrimination than a typical trichromat (55, 73) because of the interplay between additional gene copies and the coevolution of spectral tuning across rhodopsins decreases the minimal wavelength difference (Δλ) that an insect can discriminate. *Papilio* butterflies, for instance, have some of the most complex retinal mosaics known, with three ommatidial types expressing various combinations of five rhodopsin proteins (UV, B, 3 LW), as well as filtering pigments that produce diverse ommatidial spectral sensitivities (29, 45). Amino acid residues in helix 3 have been shown to mediate an absorbance shift between duplicates LW PxRh1 and PxRh3 in *Papilio (62)*, but lycaenid LW opsins possess a highly conserved helix 3 that does not exhibit any of the spectral substitutions found across Papilionidae, supporting the hypothesis that distinct spectral tuning mechanisms have evolved independently in invertebrate LW opsins. Future work will be able to disentangle the spectral tuning mechanisms of LW opsins *in vitro* using the recombinant purification system described here.

In *E. atala*, ommatidial density and expression profiling showed that photoreceptor cells containing BRh2 are less abundant (~25%) compared to those containing UV (~50%) and BRh1 (75%) rhodopsins. In spite of its low density across the eye, the derived green-shifted blue rhodopsin improves signal resolution in the green spectrum and provides direct adaptive benefits for light perception in both males and females. Compared to trichromats, the expanded six-ommatidial types primarily derive from the non-overlapping cellular expression of BRh1 and BRh2 rhodopsin mRNAs in distal rhabdomeric cells. Together, the stochastic rhodopsin mosaic offers remarkable properties that could not be achieved with the limited sensitivity of a single blue opsin gene.

In summary, our results support the scenario of a sensory system in which gene duplication generates new opsin paralogs, cis-regulatory changes generate new photoreceptor subtypes via gene-specific expression patterns, and coding mutations tune the spectral sensitivity maxima of two of the four rhodopsins (BRh2, LWRh). All of these changes have contributed to the diversification of visual spectral sensitivities, thereby enhancing colour discrimination.

### Co-evolving shifts at blue and red opsins tune intraspecific visual signalling

Additional structural features of the eye can contribute to spectral sensitivity of individual photoreceptors at long wavelengths, including lateral filtering granules found in distal rhabdomeres. However, here the primary mechanism pushing red sensitivity is through functional divergence at the LW rhodopsin locus. This remarkable novelty provides a two-factor interaction (Blue-LW) to modulate green-red sensitivity among closely related taxa.

Building on earlier butterfly microspectrophotometry work (19, 40, 41, 53), our study shows that lycaenid species with red-shifted LW opsins have their second maximum sensitivity peak and associated blue rhodopsin shifted to green maximal absorbance with λ_max_ 495-500 nm. By contrast species with a LW ancestral-green opsin with λ_max_ 520-530 nm have their second maximum sensitivity peak at blue wavelengths λ_max_ = 480-490 nm (Fig. 1C, S6A), suggesting modulatory benefits for coevolving spectral shifts at multiple rhodopsins. Within Lycaenidae, reflectance patterns of wing scales have been investigated in the genera *Celastrina* and *Callophrys* (75, 76) and these studies support a scenario of co-evolution of spectral shifts across opsins. The ventral wings visible at rest in Green Hairstreak lycaenids (genus *Callophrys*) typically appear green and consist of yellow and bluish scales with an omnidirectional reflectance peak in the green slightly above 550 nm (76). *Celastrina* species tend to display bright blue coloured wings, which predominantly reflect light in the UV-blue band (75) and can thus be perceived by comparing neuronally the spectral input from light activating retinal photoreceptors expressing both UV/ blue rhodopsins without requiring expanded red sensitivity.

These observations suggest a role for molecular tuning of blue and LW opsin genes in driving the dynamic evolution of green-red spectral sensitivity. Extant human trichromatic colour vision is similarly thought to have evolved via spectral tuning of existing short wavelength-sensitive opsins in concert with molecular variation at duplicated opsin loci leading to middle and long-wavelength sensitivities (55). Ultimately, mechanistic and functional studies of spectral tuning evolution in different orthologs in additional species will enable us to infer the ancestral states and better resolve the evolutionary time scale involved in stages of spectral tuning of blue and red sensitivities across lycaenids.

### Colour tuning improves intraspecific signal detection

Two effects of molecular variation in visual pigments in *E. atala* are that its eyes have expanded spectral sensitivity at longer wavelengths as well as increased spectral discrimination in the blue band. In order to investigate the evolutionary consequences of molecular changes in butterfly opsins in the broader context of behaviours requiring colour vision, such as finding oviposition sites, and intraspecific recognition (37, 77–80), we need to analyze the degree of overlap of spectral sensitivities (73) and evaluate this against the spectral composition of the reflected light of the background and other objects, such as that produced by foliage and conspecifics.

The limiting wavelength at which only the human L cone opsin (λ_max_ = 559-563 nm) is sensitive corresponds to approximately 625 nm (81), which is the wavelength at which orange-red can be discriminated from pure red in human colour perception. The human neighboring M cone rhodopsin is at λ_max_ = 530 nm, which at 625 nm, has a sensitivity of 0.05 (81). By extension, for *E. atala*, BRh2 has a sensitivity of 0.05 at 613 nm (Fig. 2, Fig. 6), indicating that R565 would perceive ‘pure red’ wavelengths of light from 613 nm onwards. Alternatively, we can compute the relative sensitivity of the LW rhodopsin at 700nm (3%, Fig. 1, Fig. 3), which is generally accepted as the limit for photopic vision, and then identify the corresponding wavelength at which BRh2 reaches that same sensitivity, which is 619 nm (Fig. 2). Conservatively, we can conclude that the red rhodopsin together with the green-shifted blue rhodopsin BRh2 of *E. atala* contribute to the perception of colours at long-wavelengths up to 613 nm. The LW rhodopsin sensitivity in *A. japonica* is similar to *E. atala*, and the green-red visual spectral sensitivities of *Arhopala* butterflies would capture the dorsal wing secondary reflectance peak between 500-600 nm, which is absent in *Celastrina* or *Callophrys* wings (75). Dorsal *Arhopala* wings also have velvety black edges filled with blue regions that primarily reflect light at 370 nm in males and 400 nm in females (82) and further benefit from the expanded blue sensitivity seen across lycaenid butterflies. Similarly to super-black plumage colouration in birds (83), peacock spiders (84), and recent examples in papilionids and nymphalid butterflies (85), the adjacency and sex-specific regions covered with structural velvety black scales in the lycaenid *E. atala* likely enhance the perceived brightness of nearby colour signals.

Our wing scale analysis indicates that red colour discrimination is important to see conspecifics due to their striking reflectance above 600 nm (Fig. 7A). Reflectance spectra from the red abdomen and the ventral hindwing are similar and overlap spectral sensitivities of two rhodopsins (LWRh and BRh2), meaning the butterfly can not only detect brightness but can also derive a colour signal from these patterns, and separate these input signals from those of environmental colour cues such as signals produced by foliage (Fig. 7D) (86). Abdominal reflectance in the range 613-619 nm is roughly a third of its maximal reflectance, but there is substantial overlap of the abdomen’s spectrum with BRh2 spectral sensitivity, which suggests that the Atala hairstreak’s colour perception of the abdominal colour of conspecifics is not pure red, but something more like Orange-Red. All *Eumaeus* species have this orange-red colouration and the red scale reflectance spectra are similar between the sexes, suggesting that it evolved at the base of the *Eumaeus* genera, as an aposematic signal driven by the association with cycads (87). Whereas our results show a clear sex-specific reflectance difference on dorsal wings, it is unclear whether the red warning colouration patterns may have been coopted as a conspecific signal, as in other unpalatable butterflies (79, 88), in which case it could be a by-product of the evolution of a signal used primarily to avoid predation. Hence, the L cones of most birds have red sensitivity from about 550 nm to 700 nm (59, 89), and the bright red abdominal/ hindwing reflectance in the near-infrared spectrum would stimulate mostly bird LWS /red (without MWS/ green) to perceive a colour close to “pure red”.

Lycaenid butterflies with four rhodopsins can see more colour hues in the blue-green range than mammals that have either di or trichromatic vision (34). Sex-specific scale reflectance spectra on dorsal wings convey colour information that is readily distinguishable by the visual system. Our study, by showing the expanded spectral sensitivity exhibited by lycaenid butterflies through molecular variation and functional changes in their 4-rhodopsin visual system, highlights the importance of peripheral sensory genes in driving the adaptive evolution of multi-modal communication.

## Materials and Methods

### Butterflies

Pupae of *Eumaeus atala* were collected from host plants of *Zamia integrifolia* (locally known as “coontie”) at the Montgomery Botanical Garden, Miami, FL, USA and reared at 22°C in an insectary in the MCZ laboratories under a 12:12 L:D cycle until emergence. *Callophrys sheridanii* and *Celastrina ladon* were collected on Badger Mountain, Wenatchee WA, USA and Wahclella Falls, OR, USA respectively. Eggs and young larvae of *Arhopala narathura japonica* were collected feeding on oak trees (*Quercus glauca*) from field sites near Ginoza, Okinawa, Japan and reared in the laboratory in Cambridge, MA, USA until they eclosed as adults.

### Epi-Microspectrophotometry

Quantitative epi-microspectrophotometry (epi-MSP) was used to determine absorption spectra of butterfly rhodopsins by measuring eyeshine reflectance spectra after photoconversion of the rhodopsin to its metarhodopsin product (54, 90).

Compound eyes of most adult butterfly species exhibit eyeshine, a property that allows measuring rhodopsin absorbance spectra as well as spectral sensitivity of photoreceptor pupillary responses in eyes of intact butterflies. When subjected to repeated bright white flashes under incident-light microscope, 1) there is a photochemical effect, i.e. the coloration of eyeshine changes during each flash owing to changes in absorbance spectra that accompany photo-isomerization of rhodopsins to their metarhodopsin photoproducts (90); and 2) there is a pupillary response mediated by intracellular migration of pigment granules within photoreceptor cells, causing the intensity of eyeshine to decrease rapidly with time during each flash (91).

Three MSP methods (*in vivo* Photochemistry, Optophysiology, Retinal densitometry) are critical to study rhodopsin properties. The methods are presented here, whereas the rationale upon which the three methods are based and the general procedures for using eyeshine to make photochemical measurements from butterfly eyes are described in detail in SI methods.

### *In vivo* Photochemistry

When rhodopsin is photo-isomerized to become metarhodopsin, it undergoes a spectral shift in the absorbance spectrum. For LW rhodopsins, the shift is to shorter wavelengths; the metarhodopsin peak is usually between 490 nm and 500 nm. For both UV-absorbing and blue-absorbing rhodopsins, the shift is to longer wavelengths, typically to 475 nm - 490 nm. These photochemical changes are observable in eyeshine reflectance spectra, as increased reflectance caused by loss of rhodopsin, and decreased reflectance caused by the metarhodopsin. The computed absorbance-difference spectrum (DS), therefore, has a positive peak caused by accumulation of metarhodopsin (M), and a negative peak caused by loss of rhodopsin (R) (Fig. 1, 2A).

The absorbance difference spectrum relaxes with time in the dark, but changes shape in doing so; the positive peak relaxes to zero much faster than the negative peak. The entire temporal evolution of difference spectra can be reproduced quantitatively by assuming different kinetics for the dark-processes of metarhodopsin decay and rhodopsin recovery. Metarhodopsin decay is well approximated by a single exponential process, but the time constant is a strong function of temperature. Rhodopsin recovery is considerably slower than metarhodopsin decay, making it possible to create a partial bleach using repeated episodes of bright flashes followed by dark periods during which metarhodopsin decays totally from the rhabdom. The difference spectrum for that partial bleach is a direct measurement of the absorbance spectrum of the LW rhodopsin (Fig. 1) (90). Similar experiments with photoconversion of the other spectral types of rhodopsin are more complicated because a bright blue flash designed to efficiently photo-isomerize blue-absorbing rhodopsin will also convert some LW rhodopsins to their metarhodopsins, although with less efficiency. However if the LW rhodopsins are first bleached, then difference spectra for the blue rhodopsin are measurable (Fig. 2A).

Photochemical measurements of the *E. atala* blue rhodopsin were done with a male oriented similarly to the female at Elevation 0° and Azimuth 10°. Before photoconversion of the blue rhodopsin, the LW rhodopsins were partially bleached by flashing with 20s exposure to RG645 (20s, 2s/55s). After resting in the dark for 24 min for metarhodopsins to decay, the eye was flashed with 12s RG430 (2s/60s), which converted blue rhodopsin to its metarhodopsin M505. A difference spectrum for R440 was computed from reflectance spectra measured before and 9 min after the series of bright blue flashes.

### Optophysiology

Photoreceptor cells in butterfly eyes contain intracellular pigment granules that move centripetally in response to bright illumination and deplete light from the rhabdom by scattering and absorption. This process creates an effective pupillary response observable as a decrease in eyeshine reflectance (91). This intracellular pigment migration is mediated exclusively by photo-isomerization of the rhodopsin contained within the same cell’s rhabdomere and is not influenced by physiological responses of neighboring ommatidia. Thus, the pupillary pigment granules can be used as an optically measured intracellular probe of physiological responses to light from that cell (92).

A double-beam Epi-MSP apparatus was used for optophysiological measurements of eyeshine. One beam is deep-red filtered (e.g., 710 nm), that monitors continuously the reflectance of eyeshine but does not itself cause a pupillary response. The second beam delivers a monochromatic flash that evokes a pupillary response, measured by the first beam as a decrease in deep-red eyeshine reflectance. At each stimulating wavelength, the flash intensity is adjusted with computer-controlled neutral-density wheels to produce a criterion decrease in reflectance (usually 3% - 5%). Wavelength sequence is randomized. Both flash duration and inter-stimulus interval are held constant. After completing an experimental series, the butterfly is replaced by a factory calibrated Hamamatsu S1226 photodiode and quantum flux Q measured for every criterion combination of wavelength and wheel setting. Spectral sensitivity is computed as S(λ) = 1 / Q(λ).

### Retinal Densitometry

The visual pigment content of rhabdoms can be estimated quantitatively when the tapetal reflectance spectrum is constant (white) for wavelengths shorter than about 600 nm (25, 93). A computational model based on an electron micrograph of the tracheolar tapetum shows that this “chirped” set of layers functions as a broadband reflecting interference filter exhibiting a computed reflectance greater than 90% for wavelengths between 320 and 680 nm, thereby justifying the assumption of a white reflectance spectrum in that band. This property of wideband white tapetal reflectance can be exploited to computationally estimate rhodopsin contents. It is most valuable when applied after λ_max_ of UV, blue, and LW rhodopsins have been determined from *in vivo* photochemistry (Fig. 1; Fig. 2A) and optophysiology (Fig. 2B). R360, R440, and R565 have been determined in *E. atala* (Fig. 2C-D).

The procedure is sequential. First, a reflectance spectrum was measured after all metarhodopsins had decayed from the rhabdoms, e.g., after overnight dark-adaptation (black filled circles). Next, the dark spectrum was stripped of round-trip optical density (OD) 1.50 (*male, m*) or 1.00 (*female, f*) of R565 rhodopsin so that the residual spectra (red lines) were flat from 570 nm out to the 690 nm roll-off of tapetal reflectance. Next, those residual spectra were stripped of OD 0.36 (*m*) and 0.22 (*f*) of R440 (cyan lines), leaving large dips in the blue-green around 500 nm that were poorly fit by a single blue opsin. However, stripping of OD 0.72 (*m*) and 0.55 (*f*) λ_max_ 395 (Retinal Binding Protein, RBP) left UV residues (magenta lines) well fit by density of 0.55 (*m*) and 0.55 (*f*) of R360. The remaining residuals (blue lines) were subjected to least-squares fitting to the rhodopsin template (Fig. 2E), which produced excellent R495 fits of 0.39 (*f*) and 0.70 (*m*), supporting the presence of the fourth rhodopsin, R495.

The spectral sensitivities of *Arholopa, Celastrina and Callophrys* rhodopsins were investigated using the same techniques as described in SI methods.

### *De novo* E. atala *eye transcriptome*

The heads of 10 adult males (1 day old) were dissected under ambient light with their antennae and palpi removed prior to flash freezing in liquid nitrogen. Total RNA was extracted using the Direct-zol RNA extraction kit (Zymoresearch, CA, USA). Illumina paired-end libraries were constructed using the Ultra II RNA Directional kit (New England Biolabs, USA) and sequenced with an Illumina HiSeq v4. Adaptors were removed using Trimgalore (94) and low quality reads were filtered out prior to generating a *de novo* assembly reference transcriptome for all libraries in the Trinity sequence assembly and analysis pipeline (95). This assembly resulted in 301656 transcripts with a contig N50 of 2519 bp, and average contig lengths of 1029 bp. Small fragments were filtered out based on expression values using Kallisto (96). Opsin sequences from two other lycaenids (41) were used as queries to identify all opsin mRNAs across tissues in the final assembly using BLAST (97). To confirm their identity, the candidate opsin sequences were blasted back to the NCBI non-redundant database.

### Lycaenid LW opsin cDNA characterization

RNA was extracted from eye tissue preserved in RNA shield reagent (Zymo Research) for *A. japonica, Ce. ladon and Ca. sheridanii*. Samples were first removed from the storage solution with sterile forceps, briefly blot dried on a sterile Kimwipes paper, flash frozen in a mortar containing liquid nitrogen and finely ground using a cold pestle. Following RNA purification, we quantified RNA using a Quant-iT RNA kit and a Qubit fluorometer (Invitrogen). From purified RNA, we synthesized cDNA using the The GoScript™ Reverse Transcription System (Promega) and amplified a central region of the long-wavelength opsin using degenerated oligonucleotide primers 5’- TTGAAGCTTCARTTYCCNCCNATGAAYCC-3’ (forward) and 5’-CGAATTCGTCAT RTTNCCYTCIGGNACRTA-3’ (reverse) (48). Single bands of expected sizes were obtained, and the PCR products were purified with Exo-SAP, following Sanger sequencing. We thereafter used the SMARTer RACE cDNA Amplification kit (Clontech) to prepare 5’- and 3’- RACE cDNA for each species. We carried out RACE PCRs, and to increase the specificity of RACE reactions, we performed nested PCRs for each cDNA and obtained single-band PCR products, which were gel-purified using a Qiaquick Gel Extraction Kit (Qiagen), ligated into PCR2.1 Vector kit (Invitrogen) and transformed into competent TOPO10 cells (Invitrogen). Single bacterial clones were purified, and plasmid DNAs were sequenced using M13F and M13R primers at the Harvard DF/HCC DNA Resource Core. In total we obtained sequences from 5 to 10 opsin clones for each RACE cDNA. Based on the 5’- and 3’-UTR information, gene-specific primers were designed and used in combination with respective eye cDNAs to confirm the integrity of each full-length LW opsin coding frame sequence. Opsin subfamily phylogenetic placements were confirmed by aligning selected lepidopteran opsin genes extracted from Genbank using the MAFFT package as implemented in Geneious (98), and a Neighbor-Joining (NJ) tree of the aligned dataset was constructed using RAxML (99).

### Cloning and protein expression

The coding region of each opsin transcript was amplified from eye cDNA and subcloned in a modified pFRT-TO expression vector cassette derived from pcDNA5 and containing the human cytomegalovirus (CMV) immediate early promoter (Invitrogen, USA). The expression plasmid was modified to include a C-terminal tag by the monoclonal antibody FLAG epitope sequence (DYKDDDDK), followed by a Ser-Gly-Ser linker peptide, a T2A peptide sequence (EGRGSLLTCGDVEENPG) and the fluorescent marker protein mRuby2. Plasmid DNAs were verified by Sanger sequencing and purified with the endo-free ZymoPURE™ II Plasmid Midiprep Kit (Zymo Research, USA). Two micrograms of plasmid DNAs were used to transfect small-scale HEKT293 cultures and optimize expression conditions both via mRuby2 visualization and western blot analysis. Cells were plated at a density of 0.6×10^6^ cells in a 6-well culture dish containing DMEM medium (Gibco), and transient transfection was achieved after 48h when reaching 80% confluency in a 1:3 ratio DNA (μg): PEI (μL) (Polysciences, USA) at 1mg/mL in molecular grade water, filter-sterilized at 0.22 μm. The transfected cells were harvested in cold D-PBS (Sigma-Aldrich) after 2 days, centrifuged at 4°C for 5 min at 4000 rpm, and resuspended in 50 μL Ripa lysis buffer (Invitrogen) supplemented with 1% n-Dodecyl β-D-maltoside (Sigma-Aldrich). Cell membranes were lysed for 1h at 4°C with gentle rotation on a sample homogenizer, and cell debris collected by centrifugation at 4°C for 15 min at 13,000 rpm. The crude protein lysate concentration was quantified by BSA (Sigma-Aldrich) and 25 μg crude extract was loaded on NuPAGE™ 3-8% Tris-Acetate gels (ThermoFisher) and transferred to a polyvinylidene difluoride membrane on a TurboBlotTransfer system (Biorad Laboratories). The membranes were blocked with 1% milk (Biorad) in phosphate-buffered saline containing 0.1% Tween 20 (PBS-T, Biorad) and incubated overnight with primary antibodies (aFLAG 1:2,500, aHSP90 1:50,000, GE Healthcare) containing 0.01% Sodium azide (Sigma Aldrich) on a gently rocking platform at 4°C. After washing with TPBS the membranes incubated with aFLAG and aHSP90 were respectively incubated with HRP Conjugated ECL anti-mouse and ECL anti-rabbit (Amersham, USA), revealed using the SuperSignal West Femto (Thermo Scientific) and imaged on a ChemiDoc system (Biorad Laboratories).

### Transient expression, purification of expressed rhodopsins and spectroscopy

High-expressing clones from GPCR opsin cDNAs were transiently expressed in HEKT293 cells prior to *in vitro* purification. For each construct, cells were seeded at a density of 1.0×10^6^ cells on day 0 in fifteen tissue culture dishes (10 cm diameter, ref 25382-166, VWR) in DMEM High Glucose, GlutaMAX (Life Technologies) supplemented with 10% FBS (Seradigm Premium, VWR, USA). Lipid complexes containing 24 μg DNA: 72 μL PEI (1mg/mL) diluted in Opti-MEM I Reduced Serum (Life Technologies) were added 48h later to cells reaching 75-85% confluency. Six-hours post-transfection, the culture medium was exchanged with new medium containing 5 μ.mol^-1^ 11-*cis*-retinal (2mg/mL stock in 95% Ethanol) and under dim red illumination. The *cis*-retinal absorption peak at 380 nm was confirmed using a NanoDrop™ 2000/2000c UV-VIS Spectrophotometer (Thermo Fisher) prior to each experiment using a 1:100 dilution in ethanol. Culture plates supplemented with cis-retinal were wrapped in aluminium foil and cells were incubated in the dark. Forty-eight hours post-transfection, the medium was decanted under dim red light illumination. Cells were scraped from the plates in cold filter-sterilized HEPES wash buffer (3mM MgCl2, 140mM NaCl, 50mM HEPES pH6.6-8.5 depending on protein isoelectric point) containing complete EDTA-free protein inhibitors (Sigma-Aldrich), centrifuged for 10 min at 1,620 rcf at 4°C, and resuspended in 10 mL wash buffer for two consecutive washes. After the second wash, cell pellets were gently resuspended in 10mL cold wash buffer containing 40 μM 11-*cis*-retinal. Cells expressing opsin-membrane proteins were incubated in the dark during 1h at 4°C on a nutating mixer (VWR) to increase active rhodopsin complexes, and cells were then collected by centrifugation at 21,500 rpm for 25 min at 4°C on a Sorvall WX Ultra 80 Series equipped with an AH-629 Swinging Bucket Rotor (Thermo Scientific).

Transmembrane proteins were gently extracted by pipetting in 10 mL ice-cold extraction buffer (3mM MgCl2, 140mM NaCl, 50mM HEPES, 20% Glycerol v/v, 1% n-dodecyl β-D-maltoside, complete EDTA-free protein inhibitors) and incubated for 1h at 4°C prior to centrifugation at 21,500 rpm for 25 min at 4°C. The 10mL crude extract supernatant containing solubilized rhodopsin complexes was added to 1mL Pierce™ Anti-DYKDDDDK Affinity Resin (Thermo Scientific, USA) and incubated overnight at 4°C in a 15mL falcon on a nutating mixer. Samples were loaded on Pierce™ Centrifuge Columns (ref 89897, Thermo Scientific, USA) and after 3 washes of the resin-bound FLAG-epitope rhodopsin complexes with 3-column reservoir volumes of elution buffer (3mM MgCl_2_, 140mM NaCl, 50mM HEPES, 20% Glycerol v/v, 0.1% n-dodecyl β-D-maltoside), the rhodopsin was eluted in 2mL elution buffer containing 1.25 mg (265 μM) Pierce™ 3x DYKDDDDK Peptide (Thermo Scientific, USA). The eluate was concentrated using an Amicon Ultra-2 Centrifugal Filter Unit with Ultracel-10 membrane (Millipore, USA), for 35 min at 4°C and 3,500rpm. The concentrated eluate (~350μL) was aliquoted in Amber light-sensitive tubes (VWR, USA) and kept on ice in the dark. Ultraviolet-visible absorption spectra (200-800nm) of dark-adapted purified proteins were measured in the dark from 1.5 uL aliquots using a NanoDrop™ 2000/2000c UV-VIS spectrophotometer (Thermo Fisher). Opsin purification yields were estimated following BSA analysis (Table S23, SI methods). Spectroscopic analysis was performed from the mean value of 4-6 independent spectral measurements. Raw absorbance data were fitted to a visual template (100) and polynomial functions analyses performed in R (V.0.99.486) (101)to determine the opsin maximal absorption peaks.

### Preparation of RNA probes and RNA *in situ* hybridization

We created *in vitro* transcription templates from UVRh, BRh1, BRh2, LWRh opsin complementary DNA cloned in approximately 700-base-pair (bp) segments to pCRII-TOPO (Invitrogen). Antisense cRNA probes were synthesized using T7 or Sp6 polymerases using either digoxigenin (DIG) or fluorescein (FITC) labelling mix (Sigma-Aldrich) from gel-purified PCR templates. The synthesized cRNA probes were ethanol-precipitated with NH_4_OAc 7.5M and 1 μL glycogen, spun down at 4°C for 30 min, re-dissolved in pure water and stored at −80°C. These RNA probes were first used to test mRNA expression for each opsin receptor gene. We then tested the probes by dual colour *in situ* hybridization using combinations of DIG and FITC probes to map opsin receptor expression patterns.

For *in situ* hybridization, *E. atala* compound eyes were dissected and immersed in 1.5mL eppendorf tubes containing freshly-made 4% formaldehyde (FisherScientific)/1x PBS for 2h at room temperature for fixation, then immersed successively in increasing sucrose gradient solutions (10%, 20%, 30% in PBS) for 1h each, stored in 30% sucrose solution overnight at 4°C, briefly transferred in OCT:sucrose 1:1, embedded in OCT (Tissue-Tek) and frozen in dry ice. Tangential and longitudinal eye sections (12 μm) were obtained using a cryostat (Leica), mounted on VWR Superfrost Plus Micro slides and used for RNA *in situ* hybridization following a procedure described in details previously (102). Double fluorescence *in situ* hybridization was performed using 100 μl hybridization solution (pre-hybridization buffer supplemented with 4% Dextran sulfate (Sigma) containing a combination of two opsin cRNA probes, one labeled with DIG and one labeled with FITC (at 1ng.μl^-1^ for UVRh and BRh2, and 0.5ng.μl^-1^ for BRh1 and LWRh).

### Eye anatomy

Each *E. atala* eye was immersed for prefixation in 2.5% Glutaraldehyde/2% paraformaldehyde in 0.1M Sodium Cacodylate buffer (pH 7.4) (Electron Microscopy Sciences, PA, USA) for 2h at room temperature, then stored at 4°C for 12-14h prior to fixation, embedding, ultrathin sectioning and mounting on copper grids for TEM analysis as described in SI methods.

### Homology modeling and targeted mutagenesis

To investigate the molecular basis of spectral tuning differences between the duplicated blue visual opsins, we carried out site-directed mutagenesis of amino acid substitutions that could contribute to possible changes in the maximal absorption spectra of dark-adapted rhodopsins. First, the Eat-B1 (λ_max_ = 435nm) and Eat-B2 (λ_max_ = 500nm) opsin amino acid sequences were uploaded to the SWISS-MODEL protein recognition engine (103) to generate templates aligned against the invertebrate squid rhodopsin crystal structure (PDB2Z73) (72). The predicted homology model for each blue opsin was analyzed in Pymol (104) to identify homologous binding sites in the *cis*-retinal binding pocket within a range of 5Å from any carbon in the retinal polyene chain. Of the 101 amino acid substitutions that differ between the duplicate opsins, 21 residues were predicted to interact with the *cis*-retinal chromophore, with 6 variant sites between both opsin sequences.

Amino acid sequences from blue opsins were retrieved from Genbank and aligned using the MAFFT package followed by NJ tree inference and support analysis derived from 1,500 bootstrap replicates in Geneious (97) prior to visualization in EvolView (98) with squid rhodopsin as the outgroup. The phylogeny was used to identify functionally convergent amino acid replacements repeatedly associated with similar shifts in absorption spectra between blue opsin duplicates. We identified amino acid positions that were likely to reside within the chromophore binding pocket of the opsin protein, and that also had diverging biochemical properties (charge and/or polarity).. A BRh1 plasmid DNA construct was modified to incorporate variant positions found in BRh2, namely A116S (S1), I120F (S2), G175S (S3), Y177F (S4), I206C (S5), and F207C (S6).

Chimeric BRh1 rhodopsin constructs bearing single variant sites located on helix 3 (S1, S2) and the β-strand located between helices 3 and 4 (S3, S4) were purified and analyzed by spectroscopy in the native dark state. Since only S1 and S4 had tuning effects in the range of interest, we combined coevolving adjacent site variants S1/S2, and S3/S4 and followed two distinct routes by successively adding variant sites creating triple mutants. Starting from a green-shifted BRh1 variant carrying S5 and S6, a third trajectory was studied where variant sites S1, S2, S4 were also successively added.

### Wing reflectance

Reflectance spectra were measured from leaves of *Zamia integrifolia* leaves (Zamiacae) collected at the Montgomery Botanical Garden (Miami, FL, USA), and from *E. atala* discrete wing, thorax, and abdominal patches of coloured-scales from both males and females (N_individuals_=3-5, 2 to 4 measurements per scale type, SI datafile 2), and in a Leitz Ortholux-Pol microscope equipped with a Leitz MPV-1 photometer with epi-illumination block, fitted with a Leitz 5.6X/0.15P objective. The illuminator filled the back focal plane of the objective with axial incident light. The photometer measured reflected light from the full aperture of the objective from a spot in the front focal plane that was 210 μm in diameter. Reflectance data were corrected for stray light by subtracting data measured from the MSP objective viewing a light-dump comprised of substantially out-of-focus black velvet cloth. Corrected reflectance data were normalized against the same normalization constant of 0.179 to preserve relative brightnesses among all measured body patches. Normalized reflectance data were analyzed in R (V.0.99.486) (101).

## Data accessibility

Our sequencing data have been deposited in the GenBank database under accession numbers MN831881-MN831887 and under SRA Bioproject XXXXX. Data for photochemistry and optophysiological measurements, opsin absorbance spectra, reflectance spectra and transcriptomics expression are available as supplementary files.

## Acknowledgements

We thank Masaru Hojo for providing eggs, larvae and pupae of *Arhopala japonica*, Maria Eriksson and Peg Coughin for assistance with electron microscopy, Sara Jones and Jonathan Schmidt-Burk for advice with cell culture, Ian Slaymaker for advice with protein purification procedures, Rachel Gaudet and José Velilla for guidance using Pymol and homology modeling, Rosalie Crouch for providing the 11-cis-retinal, Jeanne Serb and Davide Faggionato for advice with cis-retinal delivery, Almut Kelber, Mandyam Veerambudi Srinivasan, Mary Caswell Stoddard, Julien Ayroles, and Riannon Macrae for helpful discussions and Richard Belliveau, Maggie Starvish and Rachel Hawkins for providing logistical support. This work was supported by a Mind Brain Behavior Interfaculty grant to NEP and MAL, an Alice and Knut Wallenberg Postdoctoral fellowship at the Broad Institute of MIT and Harvard to MAL, a NSF Graduate Research Fellowship to SSa, personal funds of GDB, and NSF DEB-1541560 to NEP and PHY-1411445 to NEP and NY.

