## Supplementary material for "The evolution of red colour vision is linked to coordinated rhodopsin tuning in lycaenid butterflies": SI Appendix

**This PDF file includes:**

Supplementary text
Figures S1 to S5
Legends for Datasets S1 to S2
SI References

**Other supplementary materials for this manuscript include the following:**

Datasets S1 to S2

### Supplementary Information Text

#### Methods.

*Epi-Microspectrophotometry*

Here are some details that become important for understanding the rationale upon which the three MSP experimental methods (*In vivo* Photochemistry, Retinal Densitometry, and Optophysiology) are based. The multiple rhodopsins of a butterfly ommatidium are packaged within a long, thin rhabdom waveguide that is illuminated efficiently by cornea and cone at one end (1) and terminated optically by a tracheolar tapetum at the basal end (2). Light propagates down the rhabdom waveguide, being partially absorbed by the multiple rhodopsins and metarhodopsins as it goes. Light that survives the inward trip, and is reflected by the tapetum, propagates back up the rhabdom, again being partially absorbed. Light that survives this double pass through the rhabdom is collected by the cone and cornea, then passes out of the eye where it is observable as a narrow beam of eyeshine. Thus, the time-dependent reflectance spectrum of ommatidial eyeshine contains information about absorbance spectra of rhodopsins and metarhodopsins, reflection spectrum of the tapetum, and photoreceptor physiological responses. Eyeshine spectra also depend upon both the history of illumination and the duration of dark-time preceding measurement of an eyeshine spectrum, because the metarhodopsin content of the rhabdom decays exponentially with time in the dark, as rhodopsin content recovers at a slower rate than metarhodopsin decays.

*General procedures for using eyeshine to make photochemical measurements from butterfly eyes*

Procedures for using eyeshine to make photochemical measurements from butterfly eyes have been described previously (3-6). Briefly, a completely intact butterfly is mounted in a slotted plastic tube fixed to a goniometric stage, oriented to select a particular eye region for study. Then the microscope objective is focused inward to collapse all eyeshine into a small central spot. The field stop is adjusted to mask that spot and exclude the scattered light surrounding it. After at least two hours in the dark in warm surroundings, a reflectance spectrum is measured with a series of dim monochromatic flashes. This is used as a dark-adapted reference spectrum against which difference spectra are computed following treatment with photo-isomerizing flashes.

All LW rhodopsins shift peak absorbance to shorter wavelength upon photo-isomerization. Thus, a strategy for characterizing the spectral properties of the rhodopsin most sensitive at long wavelengths is to flash the eye using each of a graded series of sharp-cutoff, long-pass color filters, working from long-wavelength cutoff to short, measuring a reflectance spectrum after each delivery. This procedure is halted when a flash causes a measurable difference spectrum. A partial bleach is created by changing to the next filter in the series and delivering repeated flashes then monitoring dark-recovery by occasionally measuring reflectance spectra and determining the metarhodopsin decay rate. When the amount of metarhodopsin is no longer measurable, rhodopsin has not yet recovered fully so the difference spectrum, at that time, is a direct measure of the rhodopsin absorbance spectrum. An estimate of that spectrum and its $\lambda_{max}$ is obtained by non-linear least-squares fitting to Bernard's polynomial template (7).

*MSP for* Arhopala*,* Celastrina *and* Callophrys *species*

Following measurement of a dark-adapted eyeshine spectrum of *Arhopala japonica*, the eye was treated with Optics Technology interference filters, 633, 566 or 433 in separate experiments. This enabled measurement of difference spectra from partially bleached R570 (LW) and R440 (Blue) rhodopsins. Similar experiments with *Ca. sheridanii* revealed rhodopsin R518. Partial bleach of the LW rhodopsin of *Ce. ladon* could not be measured reliably owing to its narrowband blue tapetal reflectance spectrum. Optophysiological spectral sensitivities were obtained by measuring pupillary responses using 30 sec flashes. For *A. japonica*, the eye was measured at Elevation 75° and Azimuth 35º creating a log10-sensitivity function (Fig. S6A) well fit by rhodopsins R345 (UV), R440 (Blue), R500 and R570 (LW). For *Ce. ladon*, log-10 sensitivity were peaks fit by R344, R440, R489 and R520 (Fig. S6B). The absorbance spectrum for the second blue rhodopsin (R500) of *A. japonica* was computationally estimated using retinal densitometry analyses as described in the main text for *E. atala*. This was not possible for data from experiments with *Ca. sheridanii* and *Ce. ladon*.

*Eye anatomy*

Each *Eumaeus atala* eye was immersed for prefixation in 2.5% Glutaraldehyde/2% paraformaldehyde in 0.1M Sodium Cacodylate buffer (pH 7.4) (Electron Microscopy Sciences, PA, USA) for 2h at room temperature, then stored at 4°C for 12-14h. After washing with the same buffer solution at room temperature, the perfused fixed eye tissue was postfixed for 2h, washed in 0.1M Cacodylate buffer and postfixed in 1% Osmium tetroxide (OsO4)/1.5% Potassium ferrocyanide (KFeCN6), washed in water and incubated in 1% Uranyl acetate prior to successive alcohol dehydration steps. The tissue was then infiltrated in a 1:1 mixture of propylene oxide: embedding resin (TAAB Epon, Marivac Canada Inc. St. Laurent, Canada) for 48h at 4°C, prior to embedding in pure resin for polymerization at 60°C for 48h. A second resin polymerization step was performed following precise reorientation of the sample to 30° elevation in the dorsal region. Ultrathin 60 nm sections were cut on a Reichert Ultracut-S microtome equipped with a diamond blade, and transferred to a formvar/carbon coated copper grid. To increase contrast prior to examination in the microscope, the grid was incubated for 3 min in 2% Uranyl acetate in 50%

Acetone, rinsed in 50% Acetone, washed in water, and incubated in lead citrate for 30 s, blotting the excess liquid off after each incubation using a Whatman filter paper. Grids were examined on a TecnaiG2 Spirit BioTWIN (or JEOL 1200EX) electron microscope equipped with a Hamamatsu ORCA HR camera (HV=80.0kV).

*Opsin purification yields*

Large-scale transfections typically produced 15 culture dishes yielding at least 120-150.$10^6$ cells at time of harvest. Following membrane solubilization, the crude protein extract was quantified by BCA according to the manufacturer's protocol (Thermo Fisher Scientific), and then compared with the total amount of opsin protein recovered in the eluate fraction (SI Dataset 1, Table S23). On average, we quantified that the recombinant opsin proteins accounted for 216 µg /µL in 300 µL eluate, corresponding to 64.8 mg or 2.7% of the total cell protein content, although some recombinant opsins reached up to 5.5 %. This indicates that the presence of the C-terminal FLAG epitope and 20 residue-long T2A peptide chain are compatible with proper folding of monomeric opsin proteins. We further calculated the $\lambda_{280}/\lambda_{max}$ ratio for each purification, and used this as an indicator of functional rhodopsin in the protein eluate, where $\lambda_{280}$ corresponds to a combination of inactive opsins due to misfolding, mistrafficking or not forming an active rhodopsin with the chromophore in the membrane(8)(7)(7). Our procedure generally produced relatively active rhodopsins, for instance in a sample of the native Eat-LW opsin with maximal absorbance of 0.038 and an activity ratio of $\lambda_{280}/\lambda_{max}$ = 2.38.

### Supplementary Figures

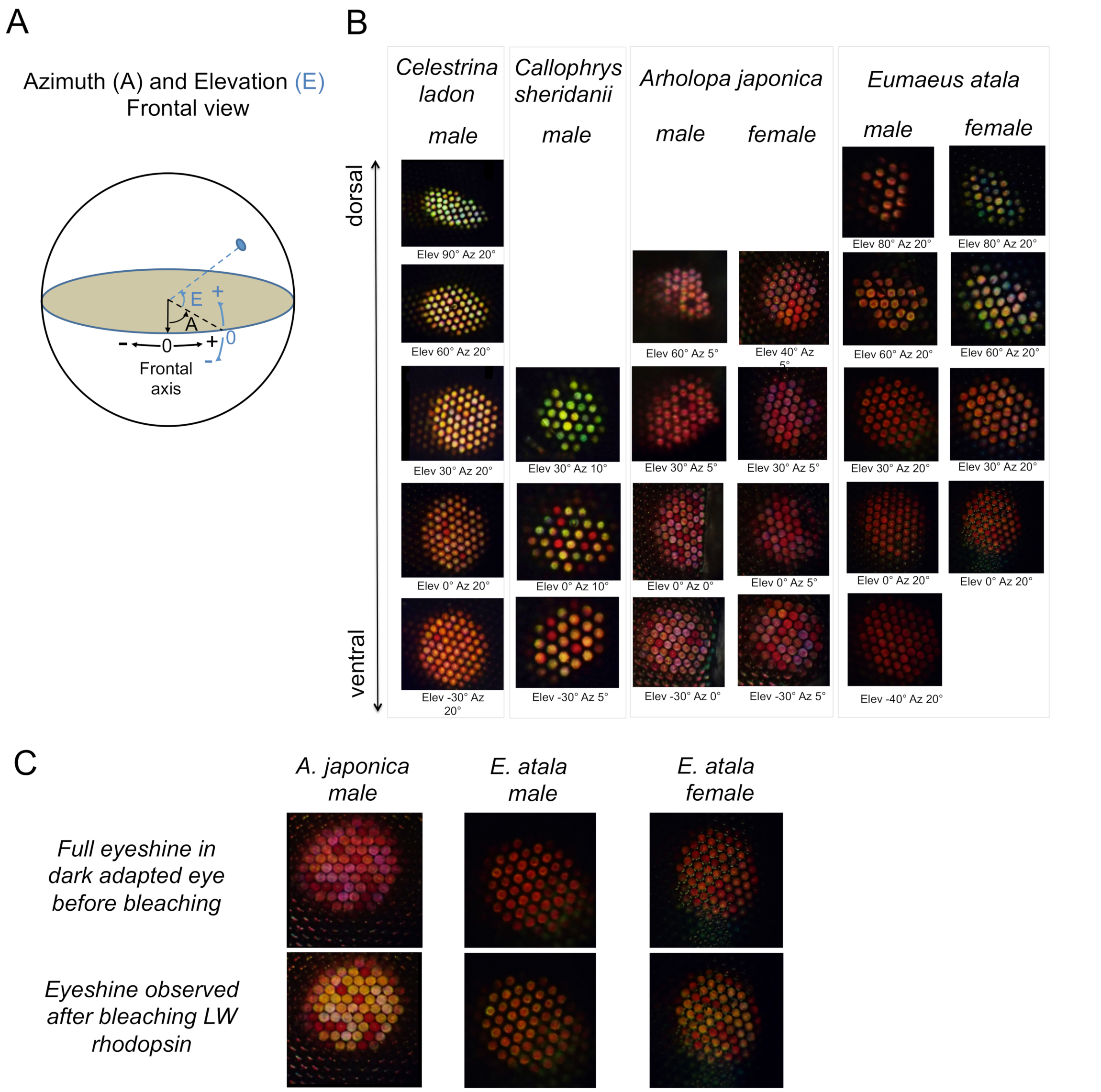


**Fig. S1. Eyeshine from surveyed lycaenid butterfly compound eyes.**

Eyeshine patterns were obtained from 21º eye patches, at various elevation angles in *Celastrina ladon*, *Callophrys sheridanii*, *Arhopala japonica* and *Eumaeus atala*. *(A)* Schematic representation of Elevation and Azimuth angles. *(B)* Epi-illumination of dark-adapted eyes shows light reflected from patches of ommatidia viewing the incident illumination from dorsal to ventral elevations. Eyeshine varies between species and also to some extent along the dorso-equatorial gradient of a single eye. In *Celastrina ladon* and *Callophrys sheridanii*, the retina is characterized by relatively homogenous green eyeshine in the dorsal part, and increasing orange eyeshine in the equatorial part of the eye, with few red ommatidia. In contrast, in *Arhopala japonica* and *Eumaeus atala*, the ommatidia appear saturated in red eyeshine, indicating the presence of red-sensitive photoreceptor cells, some of which contain filtering pigment granules located distally within the photoreceptor cells of both female and male individuals. *(C)* Change in eyeshine patterns caused by bleaching of LW rhodopsin caused by many 1 sec white-light flashes separated by 1 min dark periods, in *A. japonica* and *E. atala*. Lateral filtering pigments are not light sensitive and remain red after bleaching the LW rhodopsin.

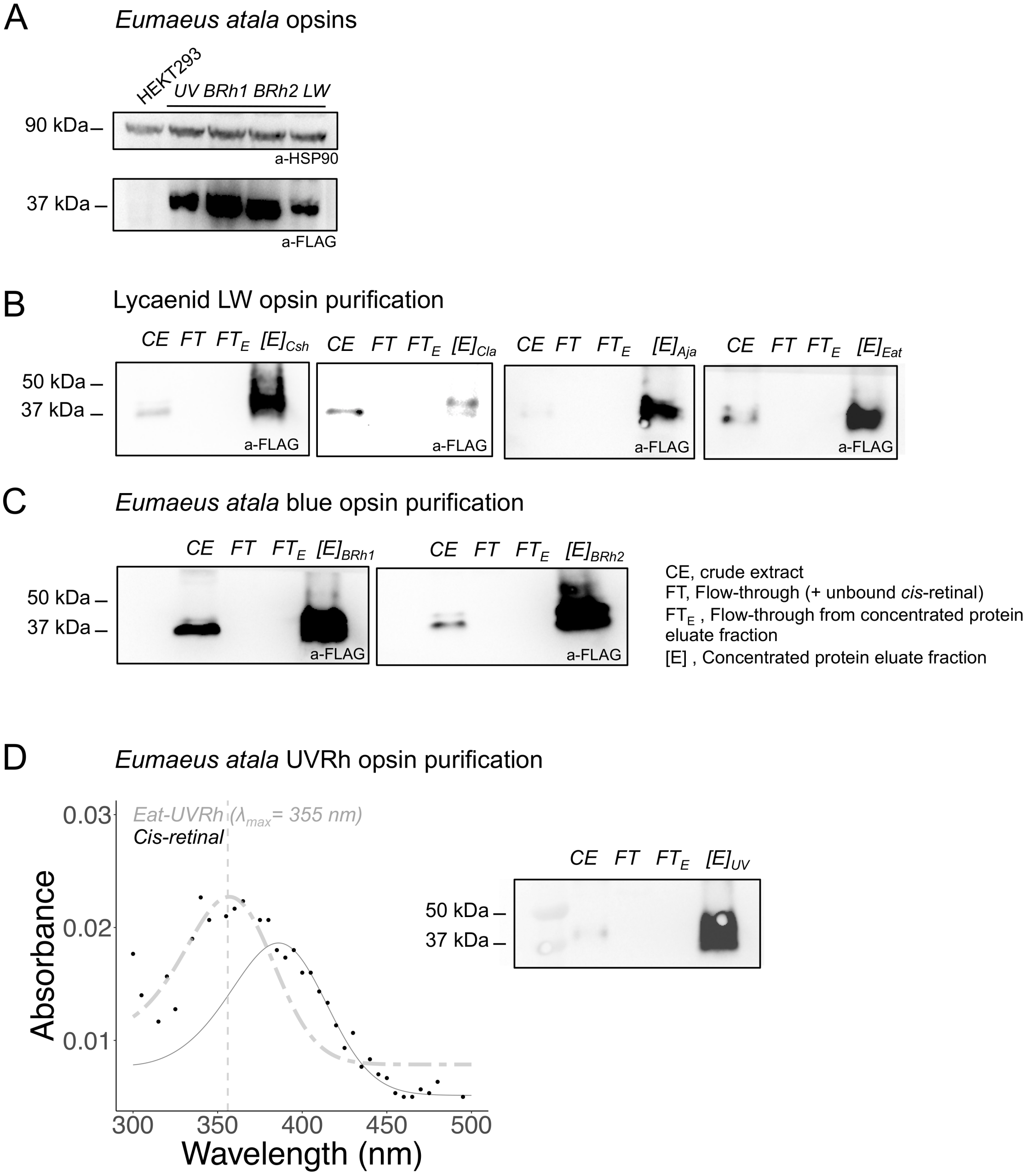


**Fig. S2. Western Blot analyses of rhodopsins reconstituted in HEKT293 cells and *E. atala* UV rhodopsin purification**

*(A)* Total membrane protein lysates of UV, Blue and LW opsins in *E. atala* following expression in the new pCDNA-FLAG-T2A-Mruby2 expression cassette (see Fig 3). HEKT293 cells were seeded at a density of $0.6.10^6$ cells in 6-well plates in DMEM medium containing 10% FBS and transfected the next day with pcDNA plasmids bearing opsin-FLAG-T2A-mRuby constructs. Transfection medium was replaced with fresh medium after 6 hours in all wells. Cells were cultured for 48 hours alongside control-untransfected control cells. Cells were washed and

harvested in cold D-PBS buffer and centrifuged for 4 min at 4,000 rpm at 4°C. The cell pellet was resuspended in Ripa extraction buffer (Invitrogen) supplemented with 1% n-dodecyl -D-maltoside (DDM) (Sigma-Aldrich) and incubated for 1 hr at 4°C, on a gently rotating platform. Debris were centrifuged at 4°C for 15 min at 14,000 rpm and the crude extract supernatant was placed on dry ice. An aliquot was subsequently used for BSA analysis. Each lane contains 20 ug total protein extract diluted in 4x Laemmli buffer (Biorad) containing 1% β-mercaptoethanol (Sigma). Samples are separated on NuPAGE 3-8% Tris-Acetate gradient gels (ThermoFisher Scientific) under non-denaturing conditions for 3 hours at 4°C and 80 Volts. After transfer to nitrocellulose membranes, low size and high size blotted bands were processed in parallel with primary antibodies anti-FLAG (1:1,000, GE Healthcare) or anti-Hsp90 (1:50,000; GE Healthcare), respectively, then with corresponding secondary antibodies ECL anti-mouse IgG (1:10,000, GE Healthcare) or polyclonal ECL anti-rabbit IgG (1:10,000; GE Healthcare), respectively. Proteins were visualized with lane 1: MW marker, lane 2: Supernatant of control nontransfected cells, lanes 3-6: supernatant of cells transfected with *E. atala* UVRh opsin (44.34 kda), BRh1 opsin (43.18 kda), BRh2 opsin (43.07 kda) and LWRh opsin (41.9 kda). The expected molecular weight of monomeric units is indicated in parentheses. *(B)* Western blot analysis of LW rhodopsin purification and *(C)* Eat-BRh1 and EatBRh2 purification. HEKT293 cells were transfected with high-expressing opsin-FLAG-T2A-mRuby constructs clones validated in a 6-well plate expression procedure. Here, transfected cells expressing rhodopsin were cultured in the dark in the presence of cis-retinal and harvested after 48hr transient expression as described in the methods. Cell membranes were solubilized and the opsin-FLAG proteins were separated from the protein crude extract using 3x-FLAG-resin column purification prior to competitive binding with FLAG-peptide The rhodopsin complexes were eluted and concentrated in Amicon-columns. CE, crude extract; FT, Column flow through; $(FT)_E$, flow through from 10-kda concentration column retaining opsin-FLAG proteins; (E), Concentrated eluate fractions. Expected molecular weight: *Celastrina ladon* LWRh, 42,11kda; *Callophrys sheridanii* LWRh, 41.94 kda; *Arhopala japonica* LWRh 41.97 kda. *(D)* Dark spectra of *Eumaeus atala* ultraviolet rhodopsin (UVRh) expressed using the HEKT293 transient cell culture system (Fig. 3A). Expected molecular weight, 44.34 kda.

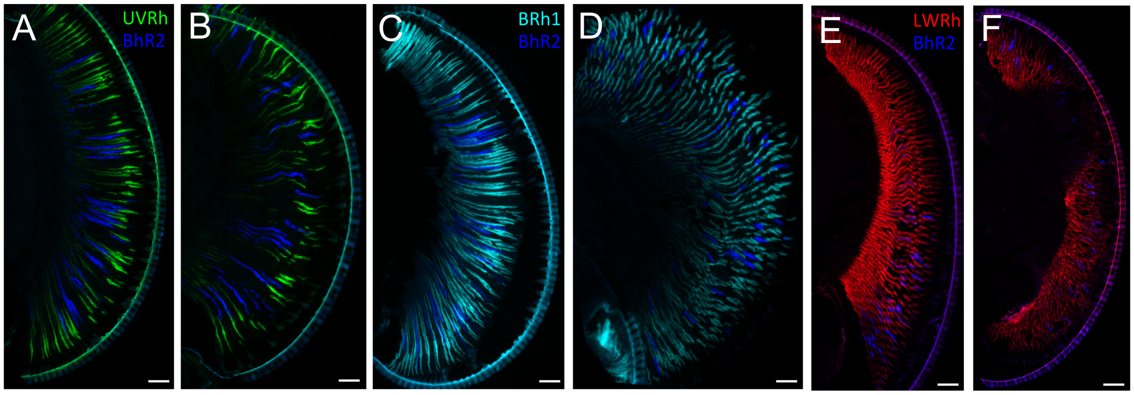


**Fig. S3**. **Sex-specific dorso-ventral expression pattern of derived BRh2 opsin duplicate**. The dorso-ventral BRh2 rhodopsin expression pattern is shown in longitudinal sections. Double fluorescent *in situ* hybridization for combinations of UVRh-BRh2 (*A,B),* BRh1-BRh2 *(C,D),* LWRh-B2Rh (*E,F*). BRh2 is expressed mainly in the equatorial and ventral female eye (*A,C,E*) and is expressed dorso-ventrally in males (*B,D,F*). Section orientation: Dorsal (top) to ventral (bottom) Scale bars, 100 µm.

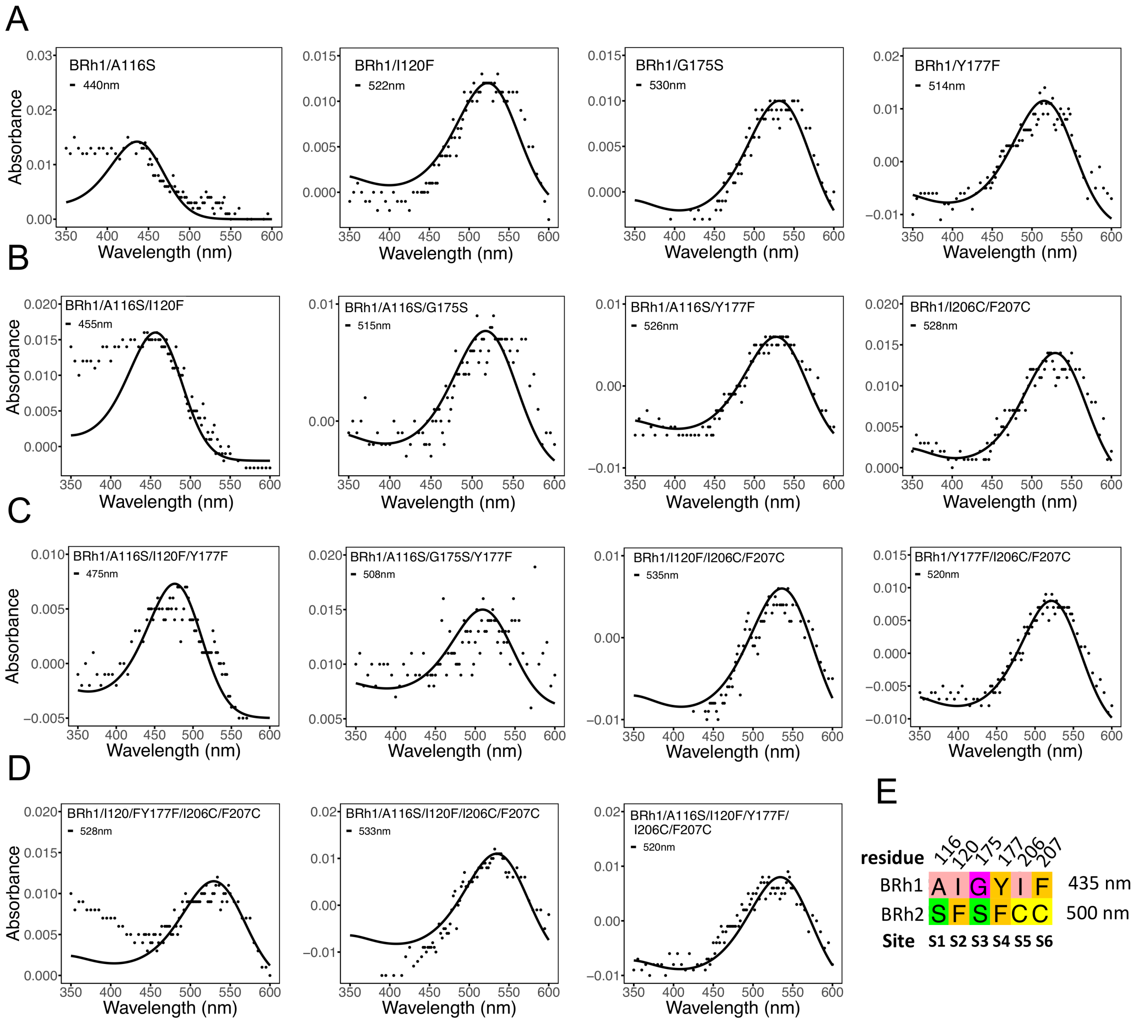


**Fig. S4. Spectral tuning effect of BRh1 rhodopsin variants.** Chimeric BRh1 rhodopsin plasmid constructs were obtained via PCR-targeted mutagenesis, expressed transiently in mammalian cell cultures and purified *in vitro* as described in the methods. Each graph represents the absorbance values for a variant rhodopsin (black dots) and the black lines represent the Bernard rhodopsin template function at maximum absorption ($\lambda_{max}$). *(A)* Single site variants, *(B)* Two-residue variants, *(C)* three-residue variants, *(D)* four and five-residue variants, *(E)* Schematic of mutated residues between blue opsin BRh1 ($\lambda_{max}$ = 435 nm) and green-shifted blue opsin BRh2 ($\lambda_{max}$ = 500 nm).

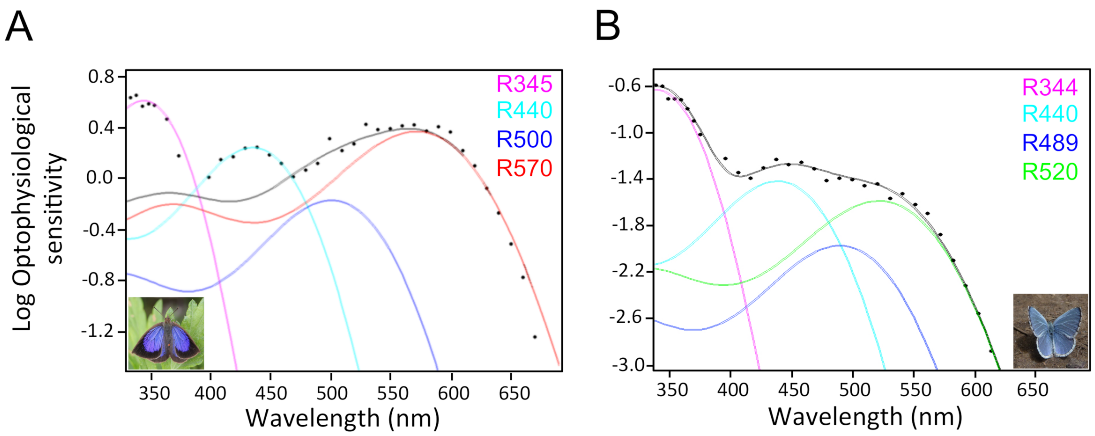


**Fig. S5. Optophysiological spectral sensitivity in lycaenid species is contributed by four rhodopsins** (magenta, ultraviolet rhodopsin; cyan, blue rhodopsin 1; blue, blue rhodopsin 2; green/red, long-wavelength rhodopsin). Black dots are optophysiological log10-sensitivity data, each point is the reciprocal of the quantum flux that actually produced the criterion pupillary response at each measured wavelength. *(A) Arhopala japonica* has long-wavelength sensitivity that expands to the far red due to a novel type of red-shifted rhodopsin receptor. *(B) Celastrina ladon* has a decreased long-wavelength spectral sensitivity conferred by a green rhodopsin receptor absorbing at shorter wavelengths.

**Dataset S1 (separate file).**

Table S1. Absorbance differences obtained by comparing a dark adapted eye with partially bleached ommatidia in *Eumaeus atala*

Table S2. Optophysiological threshold sensitivities of pupillary responses in a *Eumaeus atala* male

Table S3. Densitometric analysis of an epi-microspectrophotometric reflectance of a *Eumaeus atala* male

Table S4. Densitometric analysis of an epi-microspectrophotometric reflectance of a *Eumaeus atala* female

Table S5. Lycaenid LW rhodopsin in vitro absorbance spectra

Table S6. mRNA profiling *Eumaeus atala* opsin transcripts

Table S7. *Eumaeus atala* rhodopsin absorbance spectra

Table S8. Spectral tuning effect in Eat-BRh1 mutants

Table S9. Optophysiological sensitivity of *A. japonica* and *Ce. ladon* eyes

Table S23. Opsin purification yields

**Dataset S2 (separate file).** Contains raw reflectance data accompanying Figure 7

Table S10: *E. atala* female dorsal forewing (DFW) reflectance, cyan scales

Table S11: *E. atala* male dorsal forewing (DFW) reflectance, blue scales

Table S12: *E. atala* female ventral hindwing reflectance - black scales

Table S13: *E. atala* male ventral hindwing reflectance - black scales

Table S14: *E. atala* female ventral hindwing reflectance - cyan patch

Table S15: *E. atala* male ventral hindwing reflectance -cyan scales

Table S16: *E. atala* female ventral hindwing reflectance - red patch

Table S17: *E. atala* male ventral hindwing reflectance - red patch

Table S18: *E. atala* female abdominal reflectance

Table S19: *E. atala* male abdominal reflectance

Table S20: Reflectance from blue thorax scales in female *E. atala*

Table S21: *E. atala* male thorax blue scale reflectance

Table S22: Normalized Cycad leaf reflectance
